# The pioneer factor Zelda induces male-to-female somatic sex reversal in adult tissues

**DOI:** 10.1101/2025.03.26.645575

**Authors:** Sneh Harsh, Hsiao-Yun Liu, Pradeep K. Bhaskar, Christine Rushlow, Erika A. Bach

## Abstract

Somatic sex identity must be maintained throughout adulthood for tissue function. Adult somatic stem cells in the *Drosophila* testis (i.e., CySCs) lacking the transcription factor Chinmo are reprogrammed to their ovarian counterparts by induction of female-specific Tra^F^, but this is not mechanistically understood. Pioneer factors play central roles in direct reprogramming, and many upregulated genes in *chinmo^−/-^* CySCs contain binding sites for the pioneer factor Zelda (Zld). microRNAs repress *zld* mRNA in wild type CySCs, but they are downregulated after Chinmo loss, allowing for *zld* mRNA translation. Zld depletion from *chinmo*^−/-^ CySCs suppresses feminization, and ectopic Zld induces Tra^F^ and feminizes wild-type CySCs. *qkr58E-2* and *ecdysone receptor* (*EcR*), direct Zld targets in the embryo, are female-biased in adult gonads and upregulated in *chinmo*^−/-^ CySCs. The RNA-binding protein Qkr58E-2 produces Tra^F^, while EcR promotes female-biased gene expression. Ectopic Zld feminizes adult male adipose tissue, demonstrating that Zld can instruct female and override male identity in adult XY tissues.

**Highlights:**

- *zld* mRNA is repressed by microRNAs in XY somatic gonadal cells
- Zld is upregulated in and required for sex reversal of XY *chinmo*^−/-^ cells
- Zld induces Qkr58E-2 and EcR, which cause Tra^F^ and female-biased transcription
- Zld feminizes XY adipose cells by inducing Tra^F^ and downregulating Chinmo

## Introduction

Sex differences are observed in many adult organs during homeostasis and underlie sex-biased responses to injury and disease ^1–5^. Phenotypic differences in male and female soma arise during development as a result of several sex-biased processes, including sex chromosome constitution, dosage compensation, and steroid hormones ^6–8^. How sex differences are maintained in adult tissues is poorly understood, but gaining such knowledge is important for understanding sex-specific biological responses. The fruit fly *Drosophila* has long been an excellent model for studying sex determination during development and in adulthood ^9–11^. Work in flies has shown that sex-specific functions of adult organs such as gut and gonads are maintained by sex-biased splicing cascades, metabolism and steroid hormone responses ^1,12–15^.

In *Drosophila*, alternative splicing establishes sex-specific somatic differences based on sex chromosome constitution. In flies, XX animals are female, while XY animals are male. XX flies express the RNA-binding protein (RBP) Sex-lethal (Sxl) ^16^. Sxl binds directly to exon 2 in *transformer* (*tra*) pre-mRNA, resulting in skipping of exon 2, which contains an early stop codon, and synthesis of full-length Tra (termed Tra^F^) in females ^17,18^. XY flies lack Sxl protein, and as a result, *tra* mRNA incorporates exon 2, resulting in premature translational termination and a presumptive small peptide with no known function ^18^. In XX flies, Tra^F^ alternatively splices *doublesex* (*dsx*) pre-mRNA to produce female-specific Dsx^F^ isoform ^19–21^. As XY flies lack Tra^F^, *dsx* pre-mRNA is default-spliced and generates male-specific Dsx^M^ isoform. Dsx^F^ and Dsx^M^ are members of the DMRT family of transcription factors that regulate sex-specific differences in gene expression and external appearance in *Drosophila* ^22,23^. DMRTs have been shown to regulate sex determination across the animal kingdom ^24^.

In gonads, sex-specific somatic identity is essential for fertility and reproduction. In mammals and flies, sex-specific somatic cells are required to promote the differentiation of sex-specific gametes and to impart sex identity to germline cells ^25,26^. In both species, the production of gametes requires that the sex of the germline and the sex of the soma are the same. If somatic support cells of the gonad lose or reverse their sex identity, gonadal dysgenesis occurs with ensuing infertility ^14,15,27,28^.

*Drosophila* gonads are an excellent system to study the maintenance of somatic sex identity (**Figure S1A,B**). Both gonads have well-defined niches that support germline stem cells that ultimately give rise to gametes ^29^. Both have sex-specific somatic stem cells - cyst stem cells (CySCs) in the testis and follicle stem cells (FSCs) in the ovary - whose daughter cells encapsulate the differentiating germ cells. Somatic sex identity in male adult CySCs depends upon Dsx^M^ and Chinmo, which contains BTB and Zinc-finger domains ^14,15,30,31^. In adult ovarian FSCs, on the other hand, somatic sex identity is probably regulated by Tra^F^, as adult ovaries lacking the Tra^F^ co-factor Tra-2 degenerate, but detailed analyses were not performed ^32^. However, the direct requirement for Tra^F^ and Dsx^F^ in maintenance of adult follicle cells (FCs) has not yet been reported.

We and others have found that when CySCs in the testis lose expression of Chinmo, they lose “maleness” and become feminized (**Figure S1C**) ^14,15^. This sex reversal occurs without chromosome abnormalities and causes a catastrophic loss of spermatogenesis with ensuing infertility ^15^. We previously reported that the loss of Chinmo in CySCs causes male-to-female sex reversal by upregulating Tra^F^, which then produces Dsx^F^ instead of Dsx^M 15^. This Dsx isoform switch causes *chinmo*^−/-^ CySCs to adopt gene expression and morphology of ovarian FCs ^33^. We recently reported that these feminized somatic cells in the testis engage cell behaviors characteristic of ovarian FCs, including female-specific incomplete cytokinesis and collective rotational migration ^34^. Importantly, Sxl is not involved in *chinmo*-dependent sex reversal ^15^.

Despite this progress, we still lack mechanistic insights into how the loss of Chinmo in XY somatic cells leads to the upregulation of Tra^F^, a factor normally only expressed in XX cells^8^. Changes in gene expression and cellular behavior in *chinmo*-deficient CySCs occur within one day of knockdown, suggesting direct reprogramming of male cells into their female counterparts without an intermediate ^33,34^. Studies of direct *in vivo* reprogramming (i.e., transdifferentiation) have shown that induction of transcription factors, particularly those with pioneering activity, is a key event in this process ^35,36^.

To gain insights into whether transcription factors are induced during sex reversal, we analyzed our published transcriptomic data of *chinmo*-mutant CySCs. A significant fraction of differentially-upregulated genes in *chinmo*-mutant CySCs contain binding sites for the pioneer transcription factor Zelda (Zld) ^33^, which binds to site-specific motifs in nucleosomes and increases chromatin accessibility during zygotic genome activation in *Drosophila* ^37–42^. We show that Zld protein is low or absent in WT CySCs, despite expressing robust *zld* transcripts, but Zld is strongly upregulated during sex reversal after loss of Chinmo. These results indicate post-transcriptional regulation of *zld* mRNA in XY somatic gonadal cells. In WT CySCs, *zld* mRNA translation is repressed by microRNAs (miRs) that are present in WT male somatic cells and that are strongly downregulated upon loss of Chinmo. Ectopic Zld protein in WT CySCs can induce female-specific splicing of Tra^F^, placing Zld upstream of a key event in sex reversal and indicating that ectopic Zld can override male sex identity. Prolonged mis-expression of Zld in XY cells leads to silencing of the *chinmo* gene, demonstrating that Zld can suppress male sex identity in XY gonadal cells. We show that *qkr58E-2* and *ecdysone receptor* (*EcR*), direct targets of Zld in the embryo ^43^, are female-biased in adult somatic gonadal cells. We prove that Qkr58E-2, an RNA-binding protein, mediates *tra* pre-mRNA alternative splicing downstream of Zld. EcR has multiple critical functions in the somatic ovary ^44–48^, and the induced EcR in *chinmo*-mutant CySCs promotes female-biased gene expression. The van Doren lab showed that ectopic EcR signaling in adult XY CySCs downregulates Chinmo protein ^49^. When taken together with our results, this suggests that upregulation of EcR in Zld mis-expressing XY cells silences the *chinmo* gene. Finally, we demonstrate that ectopic Zld can feminize adult male adipose tissue. Thus, Zld can instruct female somatic sex identity and override male sex identity in two independent XY adult tissues through induction of female-biased target genes.

## Results

### Zld protein is upregulated in *chinmo*-depleted somatic cells and is required for male-to-female sex reversal

Loss of Chinmo, either through RNA interference (RNAi) or the *chinmo^ST^* allele, triggers male-to-female transdifferentiation ^14,15^. We transcriptionally profiled wild type (WT) and *chinmo*-depleted somatic cells that had been FACS-purified ^33^. Bioinformatic analyses revealed that 304 genes were significantly upregulated in *chinmo*-depleted samples ^33^. To determine whether there are common transcription factor signatures among these genes, we used iRegulon^50^. This analysis showed that 30% of upregulated genes had binding sites for Zld.

To investigate a potential role for Zld in sex reversal, we performed immunofluorescence with a Zld antibody in testes and control, tissues. As expected, Zld was expressed in nuclei of epithelial cells in the larval wing imaginal disc (**Figure S2A**) ^51^. The Zld antibody was specific, as Zld expression was lost in *dpp*-positive cells that mis-expressed a *zld-RNAi* transgene (**Figure S2B**). Zld protein was not expressed in most WT testes (**Figures 1A, C, S2C**) but occasionally, we observed faint Zld expression in a few somatic nuclei (**Figure S2D**). To further investigate Zld expression in WT testes, we used endogenously tagged alleles sfGFP-Zld ^51^ and mNeonGreen (mNG)-Zld ^52^. Both showed robust expression in larval ventral nerve cord and wing imaginal disc (**Figure S2J, K, N, O**). However, neither were expressed in WT testes (**Figure S2L, P**), supporting our results of no or very low expression of Zld in WT testes. Somatic *zld* depletion eliminated the Zld signal in all examined testes (**Figure S2E, G**), while somatic Zld mis-expression resulted in a 20-fold increase in Zld protein (**Figure S2F, H**). The faint expression of Zld in WT testes does not appear to be functionally important because depletion of Zld from the WT male somatic cells did not alter the number of CySCs (**Figure S2I**).

**Figure 1:**
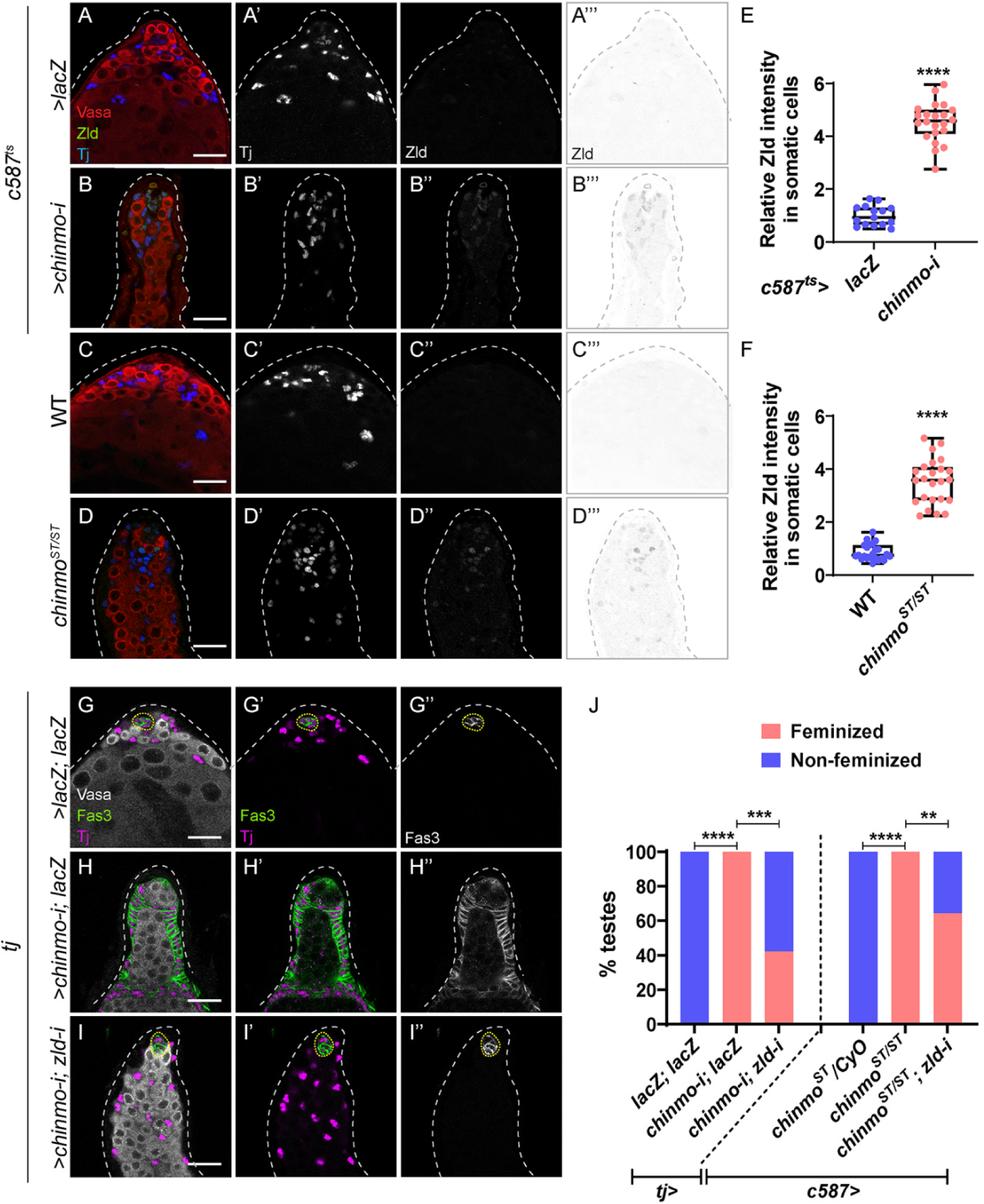
Zld is upregulated in *chinmo*-depleted somatic cells and is required for feminization. (A-D) Representative confocal images of control (*c587^ts^>lacZ*) (A), *c587^ts^>chinmo-i* (B), WT (Oregon-R) (C), and *chinmo^ST/ST^* (D) testes stained for Vasa (red), Tj (blue, grayscale) and Zld (green, grayscale, and inverted grayscale). (E, F) Graph showing relative Zld expression in the somatic cells of *c587^ts^>lacZ* (n=10) and *c587^ts^>chinmo-i* (n=14), WT (n=10) and *chinmo^ST/ST^* mutant (n=14) testes. (G-I) Representative confocal images of control testis (*tj>lacZ; lacZ*) (G), testis somatically depleted of *chinmo* (*tj>chinmo-i; lacZ*) (H) and testis with somatic co-depletion of *chinmo* and *zld* (*tj>chinmo-i; zld-i*) (I). Testes are stained for Vasa (gray), Fas3 (green, grayscale) and Tj (magenta).. (J, left side) Graph showing the percentage of feminized and non-feminized testes in *tj>lacZ; lacZ* (n=10), *tj>chinmo-i; lacZ* (n=13), and *tj>chinmo-i; zld-i* (n=45). (J, right side) Graph showing the percentage of feminized and non-feminized testes in *c587>chinmo^ST^/CyO* (n=10), *c587>chinmo^ST/ST^*(n=20) and *c587>chinmo^ST/ST^; zld-i* (n=30). Dotted lines mark the niche. Statistical analysis was performed using Student’s t-test (E, F) Fisher’s exact test (J) (** P < 0.01; *** P < 0.001; **** P < 0.0001). Scale bars: 20 µm.

Zld protein was significantly increased in *chinmo*-deficient somatic cells in *chinmo^ST^* or *chinmo*-RNAi by *c587-Gal4* in combination with thermo-sensitive inhibitor *tub-Gal80^ts^* ^53^ (termed *c587^ts^*) (3.5-fold in *chinmo^ST^* and 4.5-fold in *chinmo-RNAi*) (**Figure 1A-F**). The upregulation of Zld in feminizing testicular somatic cells raised the possibility that Zld is required for male-to-female sex reversal. To test this, we somatically co-depleted *zld* and *chinmo* and assessed the number of testes that expressed epithelial Fas3 in somatic cells after 10 days of adulthood. This assay (termed the “Fas3 assay”) has been used in past studies to measure feminization ^14,15,54^. Whereas 0% of control testes expressing two copies of a neutral *UAS* transgene (*c587^ts^>lacZ, lacZ*) had epithelial Fas3 outside of the niche, 100% of testes somatically depleted for *chinmo* (*c587^ts^>chinmo-i; lacZ*) displayed Fas3-positive, non-niche somatic cells. We were unable to test the effect of somatic co-depletion of *chinmo* and *zld* using *c587^ts^*because of high developmental lethality. Instead, we used *traffic jam* (*tj*)-*Gal4*, which is also expressed in the CySC lineage ^15^. As expected, 0% of control testes (*tj*>*lacZ, lacZ*) had epithelial Fas3 outside of the niche, and 100% of testes *tj>chinmo-i; lacZ* were feminized. Co-depletion of *chinmo* and *zld* significantly reduced feminization (42% in *tj>chinmo-i*; *zld-i* compared to 100% for *tj>chinmo-i; lacZ*) (**Figure 1G-J**). The same results were observed by somatic depletion of *zld* in *chinmo^ST^*testes (64% in *c587>chinmo^ST/ST^; zld-i* compared to 100% for *c587>chinmo^ST/ST^*) (**Figure 1J**). While *sfGFP-Zld* and *mNG-Zld* are homozygous viable, they act as hypomorphs in the adult testis as either allele significantly suppressed feminization in *chinmo-*deficient somatic cells (**Figure S2M, Q**). These results support our conclusion that Zld is required in *chinmo*-deficient cells for feminization. Upregulation of Zld in *chinmo*-mutant CySCs might reflect the normal female sex determination program in ovarian FCs, but we ruled out this possibility because Zld protein as monitored by Zld antibody or tagged alleles was not expressed in larval or adult ovary (**Figure S3A**). Zld protein was not expressed in the larval ovary, or in the larval testis, despite strong expression in the larval ventral nerve cord dissected from the same animals (**Figure S3A-D**). Furthermore, Zld depletion in adult ovarian somatic cells did not impede oogenesis (**Figure S3E**). Taken together, these results indicate that somatic loss of Chinmo significantly upregulates Zld. While Zld is dispensable for maintenance of adult male somatic cells, Zld is required for conversion of adult male gonadal somatic cells to their female counterparts.

### *zld* mRNA transcripts in WT male somatic cells are targeted by multiple miRNAs

We expected to see upregulation of *zld* mRNA in our RNA-seq analysis of *chinmo*-mutant somatic cells, but *zld* transcripts were unchanged. To validate this result, we used hybridization chain reaction fluorescent *in situ* hybridization (HCR-FISH) to quantify *zld* transcripts. As predicted, *zld* transcripts were unaltered in *chinmo*-mutant somatic cells compared with WT somatic cells (**Figure 2A-C**). These results suggest that *zld* is post-transcriptionally regulated in *chinmo*-mutant feminized cells.

**Figure 2:**
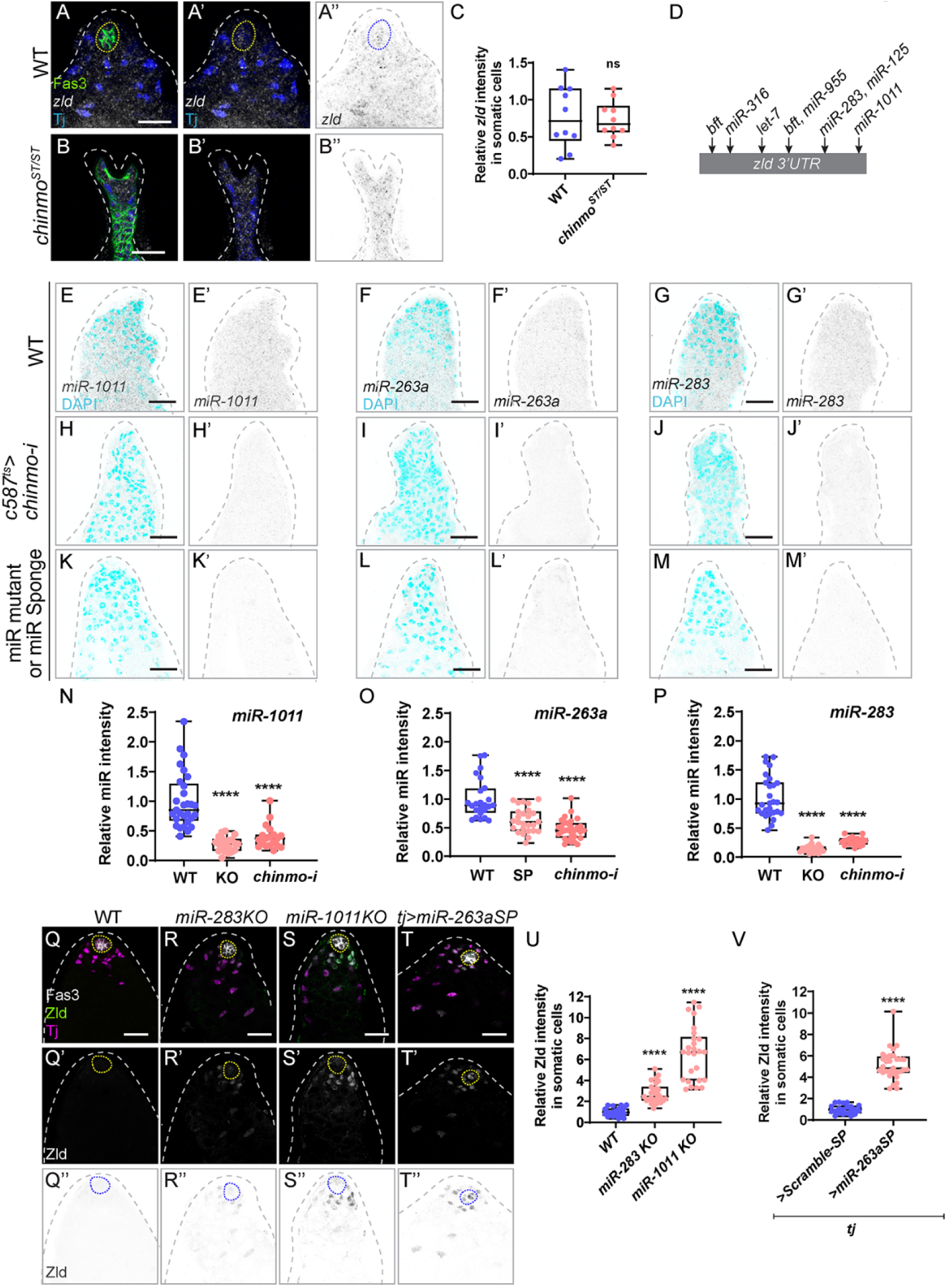
*zld* is post-transcriptionally modified in *chinmo*-deficient somatic cells. (A, B) Representative confocal images of *in situ* hybridization for *zld* mRNA in WT (Oregon-R) (A) and *chinmo^ST/ST^* testes (B). Testes are stained for Fas3 (green), Tj (blue) and *zld* mRNA (grayscale, inverted grayscale). (C) Graph showing no change in the relative *zld* mRNA intensity in the somatic cells of WT (n=8) and *chinmo^ST/ST^* feminized testes (n=10). (D) A schematic representation of *zld* 3′UTR illustrating the predicted binding sites of candidate miRNAs as identified by TargetScan. (E-M) Representative confocal images of *in situ* hybridization for *miR-1011, miR-263a,* and *miR-283* in WT (E-G), *c587^ts^>chinmo-i* (H-J) and miR-mutant or Sponge testes (K-M). miRNAs are depicted in inverted grayscale and DAPI is shown in cyan. (N-P) Graph showing relative *miR-1011* intensity in WT (n=24), *miR-1011KO* mutant (n=10) and *c587^ts^>chinmo-i* (n=13) testes (N), *miR263a* intensity in WT (n=15), *tj>miR-263aSP* (n=15) and *c587^ts^>chinmo-i* (n=13) testes (O) and *miR-283* intensity in WT (n=14), *miR-283KO* mutant (n=12) and *c587^ts^>chinmo-i* (n=8) testes (P) . (Q-T) Representative confocal images of WT (Q), *miR-283KO* mutant (R), *miR-1011KO* mutant (S), and *tj>miR-263aSP* (T) testes stained for Fas3 (gray), Tj (magenta) and Zld (green, grayscale and inverted grayscale). (U-V) Graph showing relative Zld expression in the somatic cells of WT (n=15), *283KO* mutant (n=8) and *miR-1011KO* mutant(n=10) testes (U). Graph showing relative Zld expression in the somatic cells of *tj>ScrambleSP* (Control) (n=10) and *tj>miR-263aSP* (n=12) testes (V). Dotted lines mark the niche. In Statistical analysis was performed using Student’s t-test (n.s. = not significant, P > 0.05; **** P < 0.0001). Scale bars: 20 µm.

We hypothesized that *zld* mRNA might be repressed by miRNAs in WT CySCs. To test this, we assessed Zld protein upon somatic depletion of Dicer-1 (Dcr-1), a miRNA processing enzyme. There was a significant increase in Zld protein in the *Dcr-1*-depleted somatic cells compared to WT somatic cells **(Figure S4A-C)**. We searched for potential miRNAs targeting the *zld* 3’UTR with TargetScan ^55^, which predicted seven potential miRNA binding sites in *zld 3’UTR*: two predicted sites for *bereft* (*bft*) and one predicted site each for *miR-316*, *let-7*, *miR-955*, *miR-283*, *miR-125*, and *miR-1011* (**Figure 2D**).

Based on these observations, we hypothesized that Chinmo (i.e., male sex identity) promotes the expression of one or more miRNAs that in turn repress *zld* mRNA translation in the somatic lineage. To test this model, we performed HCR-FISH using probes with initiators targeting the primary miRNA (pri-miRNA) transcripts. We then assessed which miRNAs are positively regulated by Chinmo by HCR-FISH and which miRNAs repressed Zld protein. We did not examine *let-7* since prior work had shown that it is not expressed in WT CySCs testes ^56^. HCR-FISH analysis revealed that of the remaining six miRNAs, only *miR-1011, miR-263a* and *miR-283* transcripts were present in WT testes (**Figure 2E-G, N-P**) and were significantly reduced in testis somatically depleted for *chinmo* (**Figure 2H-J, N-P**). The probes were specific to miRNA transcripts as *miR-1011, miR-263a* and *miR-283* were abolished in mutant backgrounds (**Figure 2K-M, N-P**). Zld protein was significantly increased in somatic cells in testes mutant for *miR-1011, miR-283* or *miR-263a* (**Figure 2Q-V**). These results suggest that in WT somatic cells, *zld* mRNA transcripts are silenced by at least three miRNAs whose expression is dependent on male somatic sex.

### Overexpression of *miR-263a* and *miR-1011* blocks feminization

Testes from single *miR-283, miR-1011,* or *miR-263a* mutant backgrounds did not display feminization, but they did have a significant increase in the number of CySCs, defined as Zfh1-positive, Eya-negative cells (**Figure 3A-E**) ^57,58^. Furthermore, whereas WT CySCs reside on average 25 µM from the niche, CySCs singly mutant for *miR*-*283*, -*263a*, or -*1011* reside significantly farther away (P < 0.001 for each) (**Figure 3F**). These data suggest delayed CySC differentiation in single miR mutants. These effects are specific to loss of *miR-1011* or *miR-283* and not to loss of the genes in which they are nested (*Ir93a* and *Gmap*, respectively) because somatic depletion of either Ir93a or Gmap did not affect the number of CySCs or their distance from the niche (**Figure S5** and not shown). Thus, loss of *miR-1011, miR-283*, or *miR-263a* disrupts differentiation of the somatic lineage.

**Figure 3:**
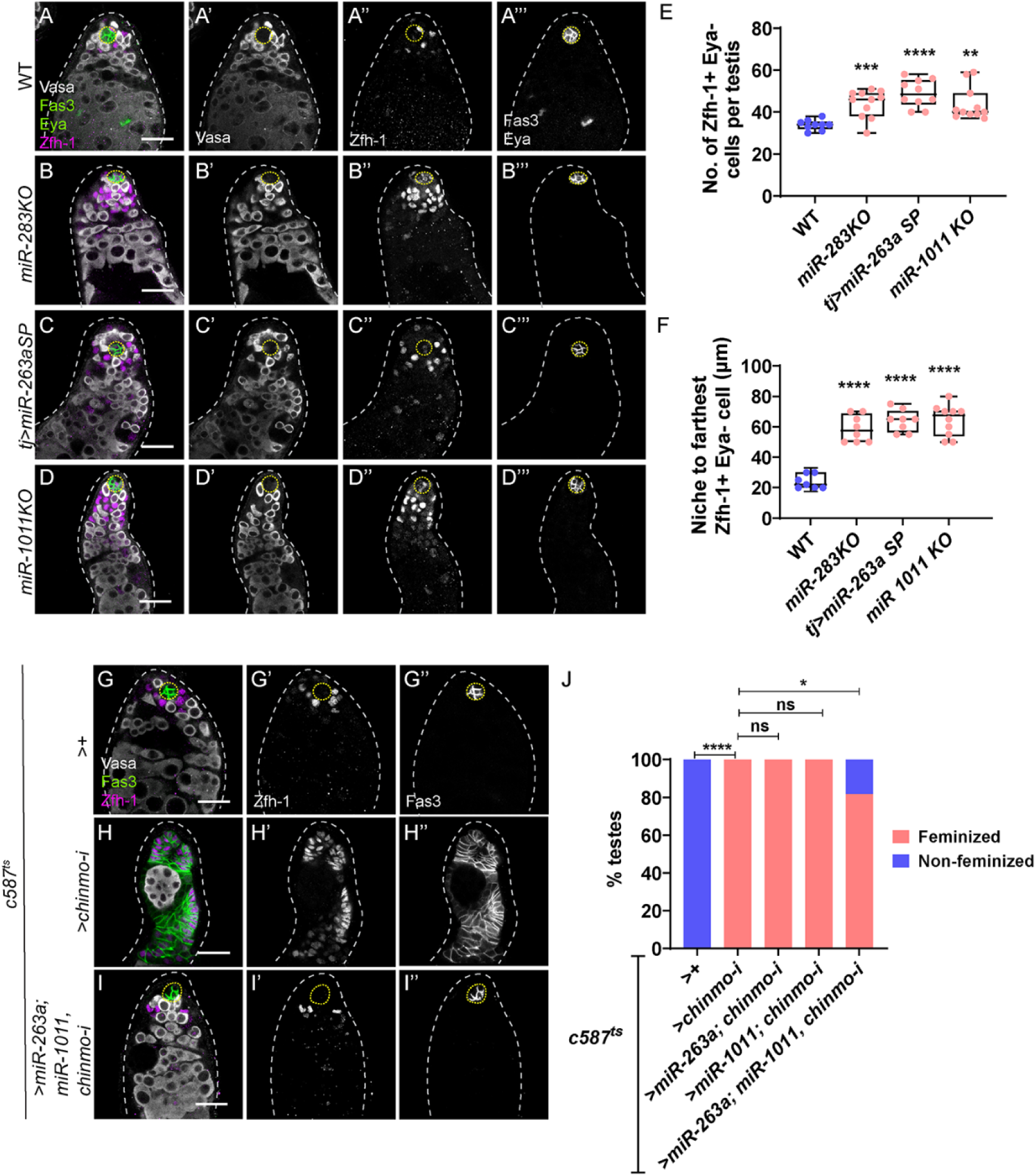
*miR-283*, *miR-1011* and *miR-263a* promote differentiation and prevent feminization of cyst cells in the testis. (A-D) Representative confocal images of WT (A), *miR-283KO* (B), *tj>miR-263aSP* (C), and *miR-1011KO* (D) testes stained for Vasa (grayscale), Zfh-1 (magenta, grayscale) and Fas3 and Eya (green, grayscale). Yellow arrow (A”) indicates a Zfh-1-positive, Eya-negative cell adjacent to the niche in WT testis. Blue arrows (B”-D”) indicates Zfh-1-positive, Eya-negative cells located many cell diameters away from the niche. The niche is outlined with a white dotted line. (E) Graph depicting the total number of Zfh-1 positive, Eya-negative cells in WT (n=9), *miR-283KO* (n=11), *tj>miR-263aSP* (n=10), and *miR-1011KO* (n=11) testes. (F) Graph showing distance of the farthest Zfh-1-positive, Eya-negative cells in WT (n=9), *miR-283KO* (n=11), *tj>miR-263aSP* (n=10), and *miR-1011KO* (n=11) testes. (G-I) Representative confocal images of *c587^ts^>+* (control) (G), *c587^ts^>chinmo-i* (H) and *c587^ts^>miR-263a; miR-1011, chinmo-i* (I) testes stained for Vasa (gray), Zfh-1 (magenta, grayscale) and Fas3 (green, grayscale). (J) Graph showing the percentage of feminized and non-feminized testes in *c587^ts^>+* (n=10)*, c587^ts^>chinmo-i* (n=20), *c587^ts^>miR-263a; chinmo-i* (n=15), *c587^ts^>miR-1011; chinmo-i* (n=18), and *c587^ts^>miR-263a; miR-1011, chinmo-i* (n=22). Dotted lines mark the niche. Statistical analysis was performed using Student’s t-test (E, F), Fisher’s exact test (J) (n.s= not significant, P > 0.05, *P < 0.05; **P < 0.01; ***P < 0.001; ****P < 0.0001). Scale bars: 20 µm.

We assessed whether increasing the level of more than one miRs could impede feminization of *chinmo*-mutant somatic cells. Co-overexpression of *miR-263a* and *miR-1011* significantly reduced the number of *chinmo*-mutant testes undergoing feminization (**Figure 3G-J**), while overexpression of these miRs individually did not (**Figure 3J**). Thus, multiple miRNAs maintain male identity by regulating somatic cell differentiation and by suppressing Zld expression, thereby preventing the activation of the female program.

### Zld is sufficient to activate female sex identity in somatic gonadal cells

We investigated whether gain of Zld is sufficient to trigger feminization by monitoring the female sex determinant, Tra^F^, using a reporter that produces GFP expression when *tra* pre-mRNA undergoes alternative splicing (*UAS-traF^ΔT2AGFP^*) (**Figure 4A**) ^15^. We previously showed that Tra^F^ is absent from WT CySCs but is present in WT ovarian FCs and in *chinmo*-mutant CySCs ^15^. To assess whether ectopic Zld could induce *tra* pre-mRNA alternative splicing, we mis-expressed the Tra^F^ sensor and *zld* in the adult somatic lineage (*c587^ts^>zld*). Flies were reared at 18°C and upshifted to the restrictive temperature of 29°C after eclosion. Tra^F^ was observed within 3–4 days of *zld* mis-expression in the adult male soma and was fully penetrant, indicating that this critical female-specific event in sex identity was occurring (**Figure 4B, C, F**). We note that ectopic Zld (*c587^ts^>zld*) induces Tra^F^ more rapidly than loss of Chinmo (*c587^ts^>chinmo-i*), which requires 7-8 days (**Figure 4D, E**), presumably because more Zld was present in the former. Tra^F^ splicing was significantly increased in Zld mis-expressing somatic cells compared to WT male somatic cells but did not reach the level of splicing in WT ovarian FCs (**Figure 4G**). Mis-expression of Zld in XY somatic cells also induced epithelial Fas3 expression outside of the niche, indicating feminization (**Figure 4B, C, H**). At 3-4 days, somatic cells with ectopic Zld still expressed Chinmo (**Figure 4I, J**, arrows), indicating that *zld* induces feminization even in the presence of male determinants. Furthermore, acquisition of Tra^F^ precedes expression of Fas3 in sex-converting somatic gonadal cells as 100% of *c587^ts^>zld* testes exhibited Tra^F^ expression while only 42% showed epithelial Fas3 expression in non-niche cells (**Figure 4H**). These results demonstrate that Zld overrides the male program and activates the female program by inducing first Tra^F^ and then Fas3.

**Figure 4:**
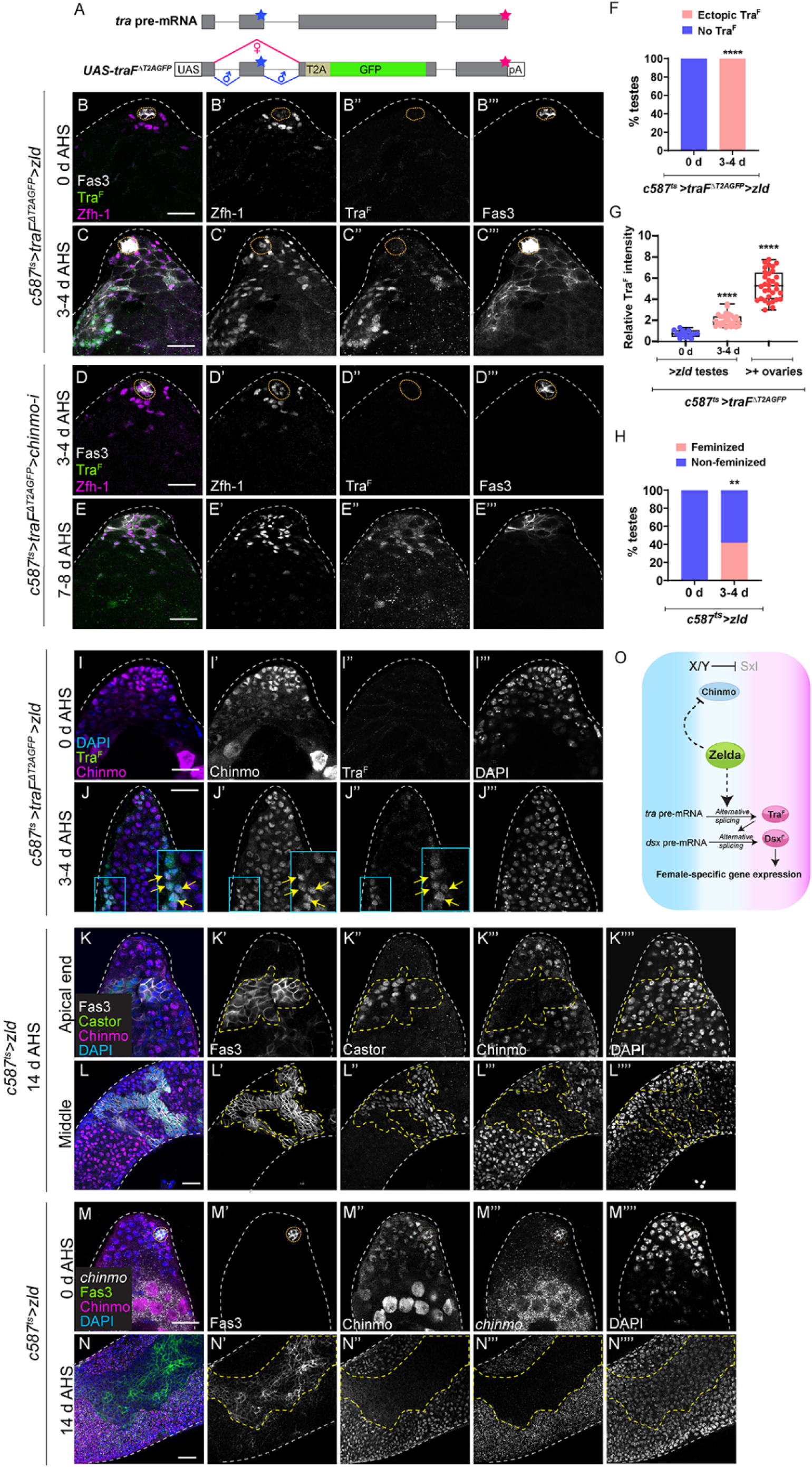
Ectopic Zld is sufficient to induce feminization. (A) Schematic of sensor for *tra* pre-mRNA alternative splicing. The transgene replaces most of *tra* pre-mRNA exon 3 with a T2A-GFP cassette and a polyadenylation signal. Gray boxes denote exons, the blue stars mark stop codon in exon 2, and the red stars mark the stop codon at the end of the sequence. Red lines indicate female-specific splicing, and blue lines show default splicing. (B, C) Representative confocal images of *c587^ts^>traF^ΔT2AGFP^*>*zld* testes at 0 days (uninduced) (A) and 3-4 days of adulthood (B). Alternative splicing of *tra* pre-mRNA (labeled Tra^F^) was monitored using anti-GFP (green, grayscale), Zfh-1 (magenta, grayscale) and Fas3 (grayscale). (D, E) Representative confocal images of *c587^ts^>traF^ΔT2AGFP^*>*chinmo-i* testes at 3-4 days (D) and 7-8 days of adulthood (E), stained for GFP (indicative of Tra^F^) (green, grayscale), Zfh-1 (magenta, grayscale) and Fas3 (grayscale). (F) Graph showing the percentage of testes exhibiting ectopic Tra^F^ (pink) and no Tra^F^ (blue) in *c587^ts^>traF^ΔT2AGFP^* >*zld* at 0 days (n=10) and 3-4 days (n=14) of adulthood. (G) Graph of the relative expression of Tra^F^ in the somatic cells of *c587^ts^>traF^ΔT2AGFP^* >*zld* testes at 0 days (n=10) and 3-4 days (n=14) of adulthood, and *c587^ts^>traF^ΔT2AGFP^>+* ovaries (n=8). (H) Graph of the percentage of feminized (Fas3-positive epithelial cells outside of the niche) and non-feminized testes in *c587^ts^>zld* at 0 days (n= 15) and 3-4 days (n=28) of adulthood. (I, J) Representative confocal images of *c587^ts^>traF^ΔT2AGFP^*>*zld* testes at 0 days (uninduced) (J) and 3-4 days of adulthood (K), stained for GFP (green, grayscale), DAPI (blue, grayscale) and Chinmo (magenta, grayscale). Arrows in J-J” in inset show somatic cells mis-expressing *zld* that induce Tra^F^ and express Chinmo. (K, L) Representative confocal images of apical (K) and middle part (L) of *c587^ts^* >*zld* testes at 14 days of adulthood stained for Fas3 (gray, grayscale), Castor (green, grayscale), Chinmo (magenta, grayscale), and DAPI (blue, grayscale). Yellow dotted lines mark feminization. (M, N) Representative confocal images of HCR-FISH for *chinmo* mRNA in *c587^ts^* >*zld* testes at 0 days (uninduced) (M) and 14 days of adulthood (N), stained for Fas3 (green, grayscale), DAPI (blue, grayscale), Chinmo (magenta, grayscale), and *chinmo* mRNA (gray, grayscale). Yellow dotted lines mark feminization. (O) Model: Ectopic Zld regulates two steps in feminization of a male somatic cell: (1) Zld triggers alternative splicing of *tra*-pre mRNA to produce Tra^F^, which activates female program; and (2) Zld suppresses Chinmo expression, which inhibits the male program. Dotted lines mark the niche. Statistical analysis was performed using Student’s t-test (G), Fisher’s exact test (F, H) (**P < 0.01; ****P < 0.0001). Scale bars: 20 µm.

### Sustained ectopic Zld suppresses male identity by silencing the *chinmo* gene

When Zld was somatically mis-expressed for 14 days, somatic cells lost Chinmo and gained Castor, a female-specific protein in gonads and a marker of feminization (**Figure 4K, L**) ^14,15^. HCR-FISH analysis revealed that prolonged ectopic Zld causes the transcriptional downregulation of the *chinmo* gene as no *chinmo* transcripts were observed in feminized (i.e., Fas3-positive) Zld-mis-expressing cells (**Figure 4N**, cells inside the dashed line), while robust *chinmo* transcripts were observed in neighboring Zld-over-expressing somatic cells that had not yet feminized (**Figure 4N**, cells outside the dashed line). These results suggest that the feminization caused by ectopic Zld occurs in two successive steps: first ectopic Zld induces the female determinant Tra^F^ that overrides the male identity program, and then Zld suppresses the male program by downregulating the *chinmo* gene (**Figure 4O**).

### Tra^F^ activation by Zld depends on Qkr58E-2, a conserved KH domain-containing RBP

Since Zld is a transcription factor, it is unlikely to participate directly in alternative splicing of *tra* pre-mRNA. Instead we reasoned that there must be an RBP that is regulated by Zld. To identify this RBP, we compared 576 annotated RBPs from the Gene List Annotation for *Drosophila* (GLAD) database ^59^, to chromatin-immunopreciptation (ChIP) targets of embryonic Zld ^43^, and to significantly upregulated genes in *chinmo*-deficient CySCs ^33^. This analysis identified Qkr58E-2, which is known to mediate alternative splicing *in vitro* ^60^. Due to a lack of existing Qkr58E-2 antibodies or protein traps, we used HCR-FISH to monitor *qkr58E-2* mRNA in adult gonads. We found that *qkr58E-2* transcripts were significantly increased (1.9 fold) in *chinmo*-mutant somatic cells (**Figure 5A-C**), consistent with our RNA-seq results ^33^. *qkr58E*-2 transcripts displayed female-biased expression in gonads with 2.5-fold higher levels in somatic ovary than the somatic testis (**Figure 5D,-E, H**). *qkr58E-2* transcripts were significantly upregulated (2-fold) in Zld-mis-expressing somatic testis at 3–4 days post-induction (**Figure 5F-H**), the time point at which Tra^F^ is robustly expressed (**Figure 4C**). We proved the specificity of two *qkr58E-2* RNAi lines because each significantly reduced *qkr58E-2* transcripts when expressed in the somatic ovary (**Figure S6A-C**). Importantly, Qkr58E-2 is required for Zld-dependent induction of Tra^F^. Tra^F^ was induced in 100% of testes somatically mis-expressing Zld but in only 40% of testes somatically mis-expressing Zld and depleting *qkr582-E* (**Figure 5I-L**). Furthermore, *qkr58E-2* is required for feminization as co-depletion of *chinmo* and *qkr58E-2* in adult male somatic cells significantly blocked feminization (**Figure 5M**).

**Figure 5:**
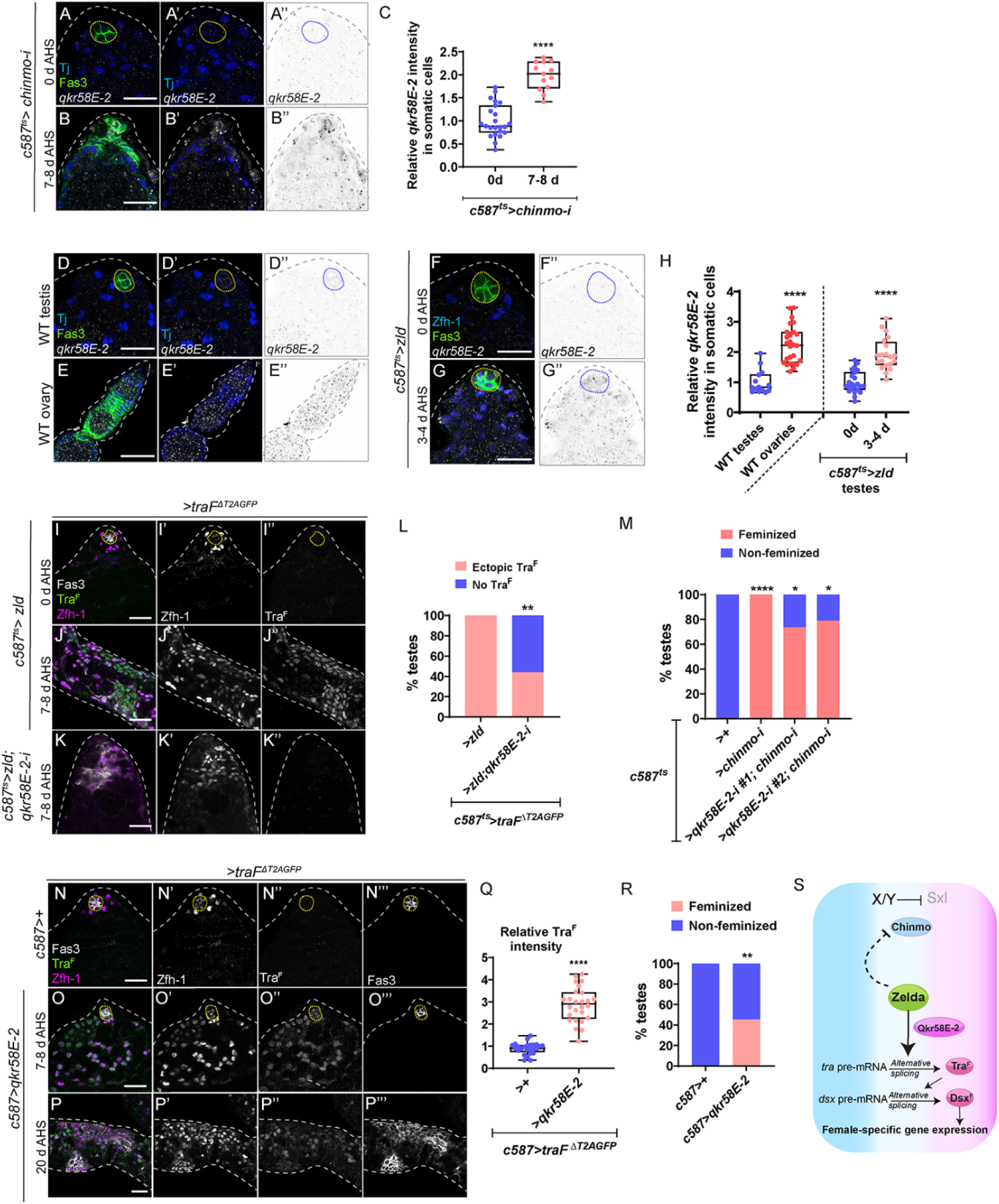
Qkr58E-2 mediates alternative splicing of *tra* downstream of Zld. (A, B) Representative confocal images of HCR-FISH for *qkr58E-2* mRNA in *c587^ts^>chinmo-i* testes at 0 days (A), and 7-8 days of adulthood (B), stained for Fas3 (green), Tj (blue) and *qkr58E-2* mRNA (grayscale, inverted grayscale). (C) Graph showing relative *qkr58E-2* mRNA intensity in the somatic cells of *c587^ts^>chinmo-i* testes at 0 days (n= 8) and 7-8 days (n= 11) of adulthood. (D-G) Representative confocal images of HCR-FISH for *qkr58E-2* mRNA in WT testis (D), WT ovary (E), *c587^ts^* >*zld* testes at 0 days (uninduced) (F), and 3-4 days of adulthood (G). The gonads are stained for Fas3 (green), Tj (blue in D, E), Zfh-1 (blue in F, G) and *qkr58E-2* mRNA (grayscale, inverted grayscale). (H) Graph showing relative *qkr58E-2* mRNA intensity in the somatic cells in: WT testes (n=8), WT ovaries (n=8) and *c587^ts^>zld* testes at 0 days (n=8) and 3-4 days (n=12) of adulthood. (I-K) Representative confocal images of *c587^ts^>traF^ΔT2AGFP^*>*zld* testes at 0 days (uninduced) (I), at 7-8 days (J), and *c587^ts^> traF^ΔT2AGFP^*>*zld; qkr58E-2-i* at 7-8 days of adulthood (K) stained for GFP (indicative of Tra^F^) (green, grayscale) and Zfh-1 (magenta, grayscale). (L) Graph showing the percentage of testes with ectopic Tra^F^ and no Tra^F^ in *c587^ts^>traF^ΔT2AGFP^*>*zld* (n=14) and *c587^ts^>traF^ΔT2AGFP^*>*zld; qkr58E-2-i* (n=11) . (M) Graph showing the percentage of feminized and non-feminized testes in *c587^ts^>+* (n=20)*, c587^ts^>chinmo-i* (n=25), *c587^ts^>qkr58E-2-i #1; chinmo-i* (n=19), *c587^ts^>qkr58E-2-i #2; chinmo-i* (n= 19). (N-P) Representative confocal images of *c587> traF^ΔT2AGFP^*>*+* (N) and *c587>traF^ΔT2AGFP^*>*qkr58E-2* testes at 7-8 days (O) and 20 days of adulthood (P) stained for GFP (indicative of Tra^F^) (green, grayscale), Zfh-1 (magenta, grayscale) and Fas3 (grayscale, grayscale). (Q) Graph showing relative Tra^F^ expression in the somatic cells of *c587> traF^ΔT2AGFP^*>*+* (n=9) and *c587>traF^ΔT2AGFP^*>*qkr58E-2* (n=10) testes. (R) Graph showing the percentage of feminized and non-feminized testes in *c587*>*+* (n=15) and *c587*>*qkr58E-2* (n=11). (S) Model: Qkr58E-2 mediates alternative splicing of *tra*-pre mRNA downstream of Zld. Dotted lines mark the niche. Statistical analysis was performed using Student’s t-test (C, H, Q), Fisher’s exact test (L, M, R) (*P < 0.05; **P < 0.01; ****P < 0.0001). Scale bars: 20 µm.

Qkr58E-2 is a bona fide regulator of *tra* pre-mRNA alternative splicing in ovarian FCs, as depleting *qkr58E-2* in ovarian somatic cells reduced both Tra^F^ and Fas3 expression in 36% of ovarioles (**Figure S7A-C**). Depletion of *qkr58E-2* in the adult somatic ovary caused severe morphological defects, including fused egg chambers (28.8%), defective germaria (18.7%), and increased cell death (52.2%), rendering females infertile (**Figure S7D, E, G**). By contrast, depletion of *qkr58E-2* in adult somatic testes had no detectable phenotype (**Figure S8A-C**). Somatic depletion of Tra^F^ target *dsx* in adult ovaries mimicked *qkr58E-2* loss (**Figure S7F, G**), highlighting the necessity of maintaining female sex identity in adulthood for proper ovary function and fertility.

### *qkr58E-2* is sufficient for feminization of XY somatic gonadal cells

To investigate whether *qkr58E-2* is sufficient to induce Tra^F^ expression, we misexpressed *qkr58E-2* using *qkr58E-2^G^*^3095^, a P-element containing *UAS* sites inserted in the *5’UTR* of *qkr58E-2*. Indeed, *qkr58E-2* transcripts were increased 6-fold in *c587>qkr58E-2^G^*^3095^ (**Figure S6D-F**). Somatic *qkr58E-2^G^*^3095^ misexpression caused a significant 2.9-fold increase in Tra^F^ compared to WT male somatic cells (**Figure 5N, O, Q**). Furthermore, somatic Qkr58E-2 mis-expression induced feminization in 45% of testes (**Figure 5P, R**). These results suggest that Qkr58E-2 is female-biased and is a key mediator of *tra* pre-mRNA alternative splicing in *chinmo*-mutant and Zld-misexpressing somatic gonadal cells (**Figure 5S**).

### Ectopic EcR expression is an early event in *zld*-mediated male-to-female sex reversal

We investigated how prolonged misexpression of Zld silences the male program by downregulating Chinmo expression. We first tested whether upregulation of female determinants was sufficient to repress Chinmo. However, prolonged misexpression of Tra^F^ or Dsx^F^ in the somatic testis for 20 days did not affect Chinmo expression or induce feminization (**Figure S9A-D**), consistent with previous reports ^14,15^.

We next considered EcR as a candidate because it is a direct Zld target in the embryo ^43^, *EcR* mRNA is upregulated in *chinmo*-mutant CySCs ^33^, and EcR causes downregulation of Chinmo in other *Drosophila* tissues ^61^. Additionally, the van Doren lab reported that EcR protein exhibits female-biased expression in larval gonads and that mis-expression of EcR in adult somatic testis causes loss of Chinmo protein and their subsequent feminization ^49^. Using the Ag10.2 antibody that recognizes all EcR isoforms, we found that EcR was present in WT adult ovarian FCs. This expression was lost when *EcR-RNAi* was expressed in adult FCs (*c587^ts^>EcR-i*), confirming antibody specificity (**Figure 6A, B, E**). Using the same antibody master mix and the same confocal settings, we found EcR was absent from CySCs and early cyst cells but was occasionally observed in late cyst cells in WT testes (**Figure 6C**, arrows). Indeed, EcR levels were 3-fold higher in the somatic ovary than testis (**Figure 6E**). Consistent with upregulation of *EcR* mRNA in *chinmo*-mutant CySCs, EcR protein was significantly upregulated in these cells (**Figure 6D, E**). Furthermore, known EcR target genes were significantly increased in *chinmo*-mutant CySCs, including *Hr3* (30-fold P^adj^ < 0.034) and *Eip63F-1* (5.4-fold P^adj^ < 0.095), indicating that EcR signaling occurs in *chinmo*-mutant CySCs.

**Figure 6:**
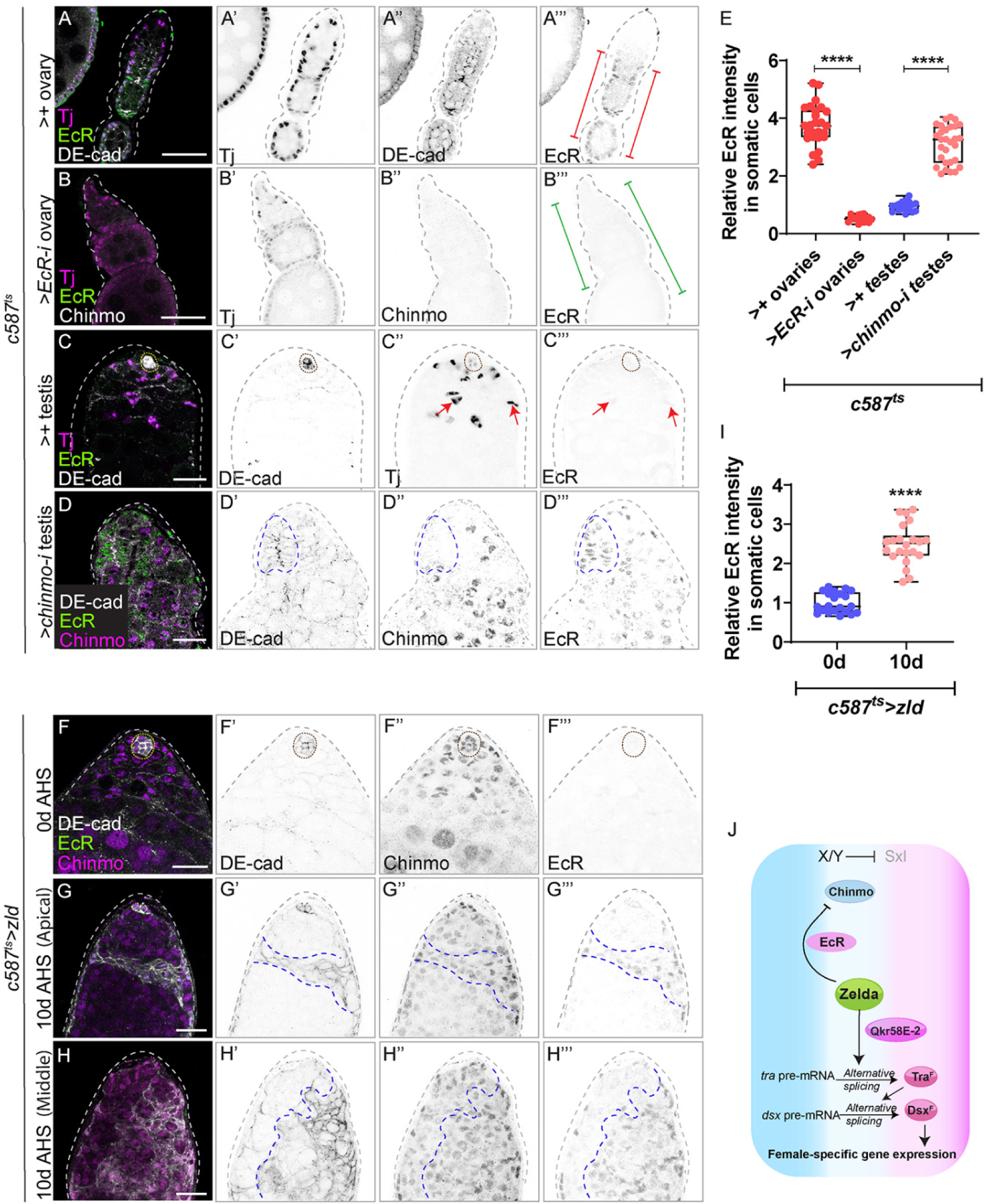
*zld* mis-expression in male somatic gonad triggers EcR expression. (A-D) Representative confocal images of *c587^ts^>+* ovary (A), *c587^ts^* >*EcR-i* ovary (B), *c587^ts^>+* testis (C), and *c587^ts^* >*chinmo-i* testis stained for DE-cadherin (gray, inverted grayscale), EcR (green, inverted grayscale), Tj (magenta, inverted grayscale) and Chinmo (magenta, inverted grayscale). Red brackets in A”’ show robust EcR expression in the ovarian follicle cells. Green brackets in B”’ show diminished expression of EcR in the follicle cells. Red arrows in C” and C”’ show faint EcR expression in the late differentiated cyst cells. Blue dashed line in D’-D’” D”’ show ectopic EcR expression in *chinmo*-depleted somatic cells. (E) Graph showing relative EcR intensity in the somatic cells in: *c587^ts^>+* ovaries (n=8), *c587^ts^>EcR-i* ovaries (n=12), *c587^ts^>+* testes (n=8), and *c587^ts^>chinmo-i* testes (n=8). (F-H) Representative confocal images of *c587^ts^>zld* testes at 0 days (uninduced) (F) and apical (G) and middle part (H) of *c587^ts^>zld* testes at 10days of adulthood stained for DE-cadherin (gray, inverted grayscale), Chinmo (magenta, inverted grayscale) and EcR (green, inverted grayscale). Purple dotted lines in G’, G” and G”’ and H’, H” and H”’ mark feminized cells. (I) Graph showing relative EcR intensity in the somatic cells in *c587^ts^>zld* testes at 0 days (n=8) and 10days (n=10) of adulthood. (J) Model: EcR represses the chinmo gene downstream of Zld Dotted lines mark the niche. Statistical analysis was performed using Student’s t-test (****P < 0.0001). Scale bars: 20 µm.

We tested whether *zld* misexpression induces EcR expression. Because *chinmo* loss also induces EcR (**Figure 6D, E**), we analyzed EcR expression at a time point when Chinmo was still present (i.e., 10 days of *zld* misexpression). Ectopic Zld significantly increased EcR expression in most somatic cells (**Figure 6F-H)**). As expected, these EcR-positive cells also expressed Chinmo (**Figure 6G”, H”**). These results indicate that ectopic EcR, similar to Qkr and Tra^F^, is an early event in male-to-female sex reversal (**Figure 6J**).

### Ectopic Zld feminizes adult male fat body

We next wanted to test whether ectopic Zld could cause sex reversal in other adult male tissues. We chose the adult fat body (i.e., *Drosophila* adipose tissue), which comprises sexually dimorphic polyploid cells. We found that male adipose cells expressed Chinmo and Dsx^M^ (Dsx^M^::GFP), while female cells expressed Tra^F^ and the Dsx^F^ target Yolk protein 1 (Yp1, Yp1::GFP) ^22,62,63^ (**Figures 7A-F, S10**). Male adult fat cells mis-expressing Zld induced alternative-splicing of *tra* pre-mRNA (**Figure 7G-I**) and significantly downregulated Chinmo protein (**Figure 7G, J**). Thus, ectopic Zld can induce female sex identity program and repress male identity in two somatic tissues, gonad and fat body.

**Figure 7:**
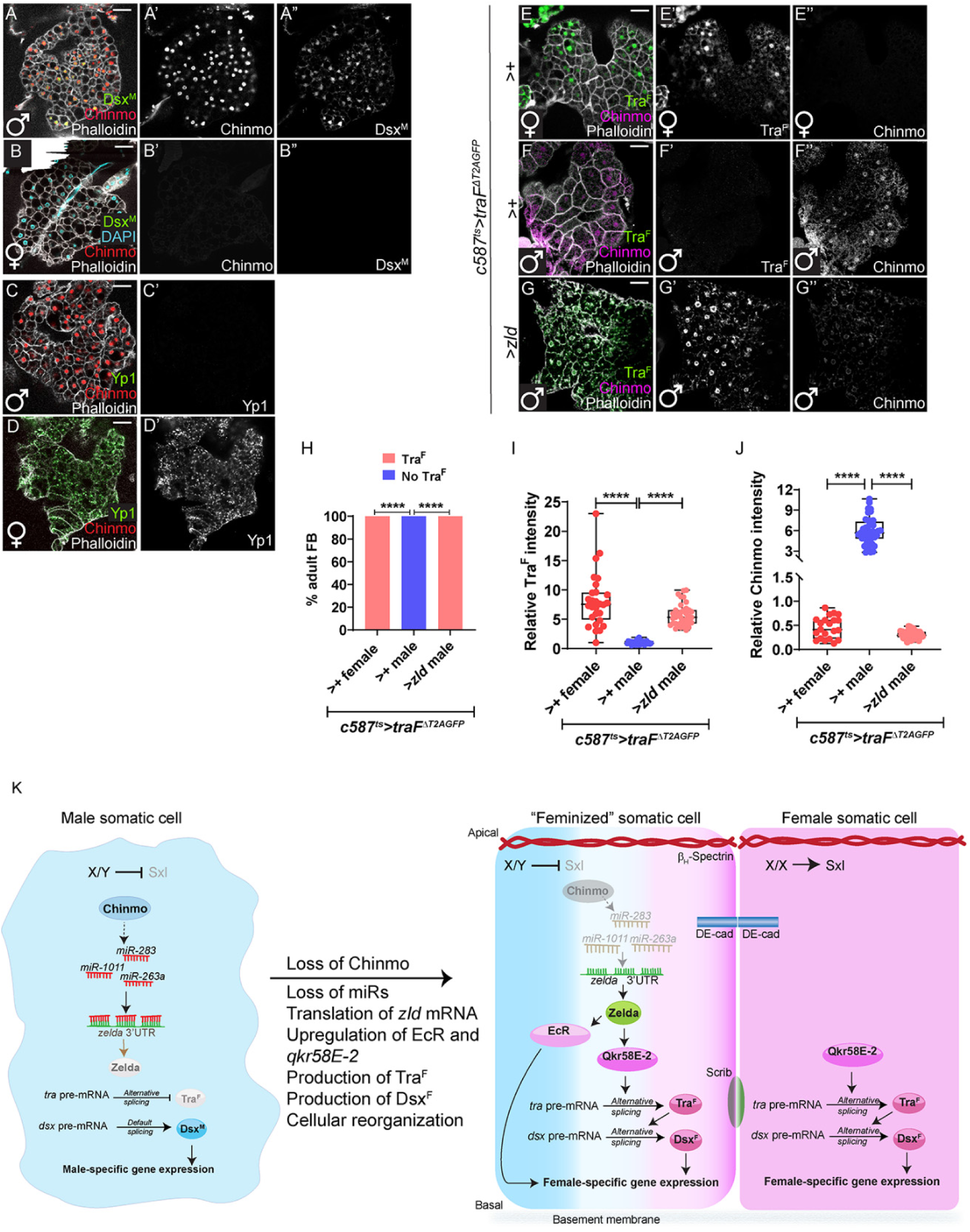
Ectopic *zld* feminizes male adipose tissue. (A, B) Representative confocal images of male (A) and female (B) Dsx^M^::GFP adult fat body stained for Chinmo (red, grayscale), Phalloidin (grayscale) and Dsx^M^ (GFP, green, grayscale). (C, D) Representative confocal images of male (C) and female (D) Yp1::GFP adult fat body stained for Chinmo (red, grayscale), Phalloidin (grayscale) and Yp1 (GFP, green, grayscale). (E-G) Representative confocal images of *c587^ts^>traF^ΔT2AGFP^*>*+* female (E), *c587^ts^> traF^ΔT2AGFP^*>*+* male (F), and *c587^ts^>traF^ΔT2AGFP^*>*zld* male fat body (G) stained for GFP (indicative of Tra^F^) (green, grayscale), Phalloidin (grayscale) and Chinmo (magenta, grayscale). (H) Graph showing the percentage of testes exhibiting ectopic Tra^F^ (pink) and no Tra^F^ (blue) in *c587^ts^>traF^ΔT2AGFP^*>*+* female (n=7), *c587^ts^>traF^ΔT2AGFP^*>*+* male (n=8), and *c587^ts^>traF^ΔT2AGFP^*>*zld* male fat body (n=12). (I) Graph showing relative Tra^F^ expression in *c587^ts^>traF^ΔT2AGFP^*>*+* female (n=7), *c587^ts^>traF^ΔT2AGFP^*>*+* male (n=8), and *c587^ts^>traF^ΔT2AGFP^*>*zld* male fat body (n=12). (J) Graph showing relative Chinmo expression in *c587^ts^>traF^ΔT2AGFP^*>*+* female (n=7), *c587^ts^>traF^ΔT2AGFP^*>*+* male (n=8), and *c587^ts^>traF^ΔT2AGFP^*>*zld* male fat body (n=12). (K) Model. (Left) In male somatic gonadal cells, Chinmo positively regulates expression of *miR-283, miR-263a,* and *miR-1011*, which suppress *zld* mRNA, thereby maintaining low Zld protein levels and enforcing the male program. (Middle) Loss of Chinmo disrupts this regulatory network, leading to reduced miRNA expression and translation of *zld* mRNA. Elevated Zld initiates two key feminization processes: (1) activation of *Qkr58E-2*, which facilitates alternative splicing of female-specific *tra* pre-mRNA to Tra^F^, triggering the female program, and (2) upregulation of the EcR, a crucial regulator of follicle cell development and oogenesis. Statistical analysis was performed using Fisher’s exact test (H) and Student’s t-test (I, J) (****P < 0.0001). Scale bars: 20 µm.

## Discussion

The key findings of this study (**Figure 7K**) are that (1) Zld is repressed by miRs in WT male CySCs but is translated upon loss of Chinmo; (2) gain of Zld in *chinmo*-mutant CySCs leads to upregulation of Qkr58E-2 and EcR, both of which are female-biased in the somatic gonad; (3) Qkr58E-2 produces Tra^F^, which in turn generates Dsx^F^; (4) EcR promotes female-biased gene expression by upregulating targets like *Hr3* and *Eip63F-1*. Thus, Zld lies upstream of the production of two female-biased transcription factors - Dsx^F^ and EcR - that serve critical roles in the ovary. We further showed that ectopic expression of Zld in the WT male somatic gonad causes feminization through Qkr58E-2 and EcR. Since ectopic EcR signaling in WT male soma causes Chinmo protein loss and feminization ^49^, we favor the interpretation that the induction of EcR in Zld mis-expressing CySCs leads to the downregulation of the *chinmo* gene. Finally, we show that ectopic Zld converts adult male adipose tissue to its female counterpart.

Our work demonstrates a critical role for Qkr58E-2 in maintaining female sex identity in adult ovarian FCs. Somatic loss of Qkr58E-2 from adult ovaries leads to defective oogenesis. Interestingly, we show that adult somatic depletion of Dsx^F^ yields similar ovary phenotypes; this is, to the best of our knowledge, the first demonstration of the requirement for Dsx in the adult somatic ovary. Collectively, these results indicate that somatic female sex identity needs to be maintained in the adult ovary, similar to the need for sustaining somatic male sex identity in the testis ^14,15^. Intriguingly, KHDRBS1, the mouse homolog of Qkr58E-2, is required somatically for female fertility in mice. Since human patients with missense mutations in *KHDRBS1* have premature ovarian failure ^64–67^, it is likely that our work has direct relevance to human fertility.

This study demonstrates that EcR is female-biased in adult somatic gonadal cells. EcR is strongly expressed in ovarian FCs. We observe EcR occasionally in differentiating somatic cells in the testis but not in CySCs. This is in contrast to prior work, which reported EcR expression throughout the testis somatic lineage ^68,69^. We are at a loss to reconcile these discrepancies, but we are confident with our results because *EcR* depletion from FCs significantly reduced EcR (**Figure 6A, B, E**). We used the same EcR antibody master mix to stain testes and the same confocal settings to image EcR expression. Our data suggest that EcR signaling is absent or very low in WT testis soma, which is consistent with prior reports showing that EcR is dispensable in WT testes ^44,69^. Our model is also consistent with recent work demonstrating that mating induces EcR signaling in CySCs, which then disrupts somatic encystment of male germ cells ^70^. Thus, suppressing EcR signaling in the adult male somatic gonad is important for spermatogenesis.

The fact that EcR signaling can repress Chinmo in adult somatic testis raises the question of whether the lack of Chinmo in the adult somatic ovary is due to the presence of EcR in these cells. Our results suggest that this is not the case since Chinmo is still absent in the adult follicle cells depleted of EcR (**Figure 6B”**). However, it is possible EcR represses Chinmo during ovary development, and this repression is maintained in adulthood. We do not understand how ectopic Zld represses *chinmo* in the adult somatic testis. In larval neuroepithelium, EcR silences the *chinmo* gene through an unknown mechanism ^61^. In the larval wing disc, downregulation of *chinmo* correlated with an increase in EcR, but EcR did not bind the *chinmo* locus, suggesting an indirect effect ^71^. Chinmo and the EcR target Broad (Br) are reciprocal antagonists during development, possibly at the level of transcription ^72,73^, but adult CySCs already express Br ^69,70^, so this mechanism is unlikely. Future work using CUT&RUN and ATAC-seq will be needed to determine how EcR represses the *chinmo* gene in male somatic cells.

Our work shows that Zld protein is not expressed in the third instar larval or adult ovary. This result was unexpected because Zld promotes female sex identity in the early embryo by promoting expression of the *Sxl* “establishment” promoter that is required for the first burst of Sxl, which then auto-regulates its own expression through the Sxl “maintenance” promoter ^16^. This suggests that Zld is required very early in development to induce female somatic sex but is not needed later for sex identity maintenance. Interestingly, the Nystul lab reported that *zld* mRNA was among the top 50 transcripts in FSCs in the adult ovary ^74^. Taken together with our data, this suggests that *zld* mRNA may be latent in adult somatic gonad cells, perhaps as a result of their shared development. The reason for this latency is not yet clear. Future work will be needed to determine whether *zld* mRNA is repressed in other adult somatic cells and whether this occurs through *miR-1011*, *-283*, and *-263a*. We have shown that ectopic Zld can feminize two *dsx*-positive somatic tissues. However, recent work has shown that Tra^F^ is functional in all female cells, regardless of *dsx* status ^8^. Therefore, it will be important to determine whether ectopic Zld can feminize *dsx*-negative cells, which comprise the majority of cells in the female fly. Finally, follow-up studies will be needed to determine whether Zld acts as a pioneer factor in adult male cells in the somatic gonad and adipose tissues. Addressing this will require ATAC-seq methodologies.

### Limitations of the study

Despite the fact that both *qkr58E*-2 and EcR increase in *chinmo*^−/-^ CySC, we have not formally proven that they are direct targets of Zld as they are in the embryo. Our results suggest that Qkr58E-2 protein increases in *chinmo*^−/-^ CySCs but we have not demonstrated this because of the lack of antibodies or endogenously-tagged alleles. Our results strongly suggest that *zld* mRNA is repressed by *miR-1011*, *-283*, and *-263a*, but we cannot rule out the possibility that other miRs are not involved.

## TAR METHODS

### Key Resources Table

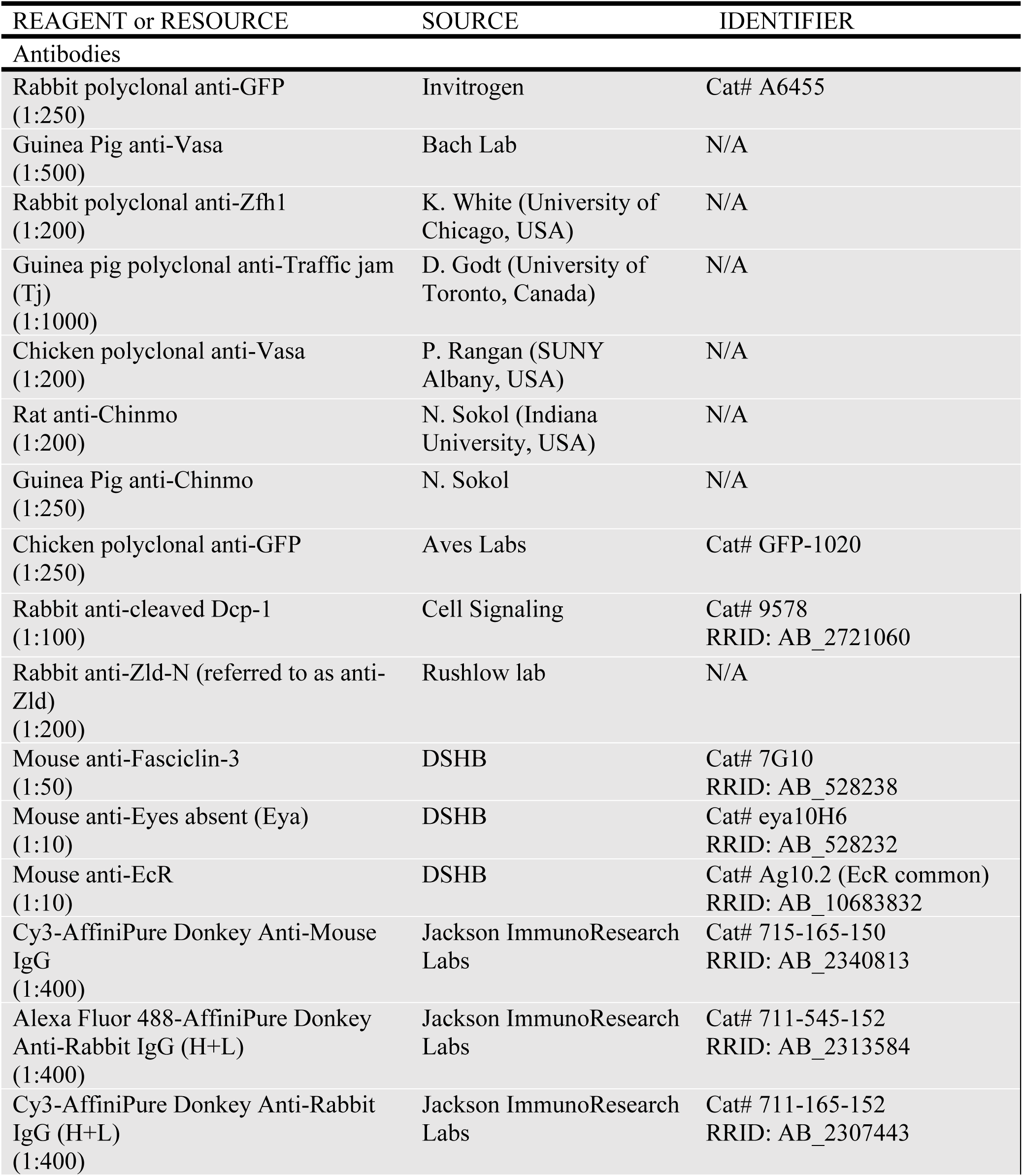
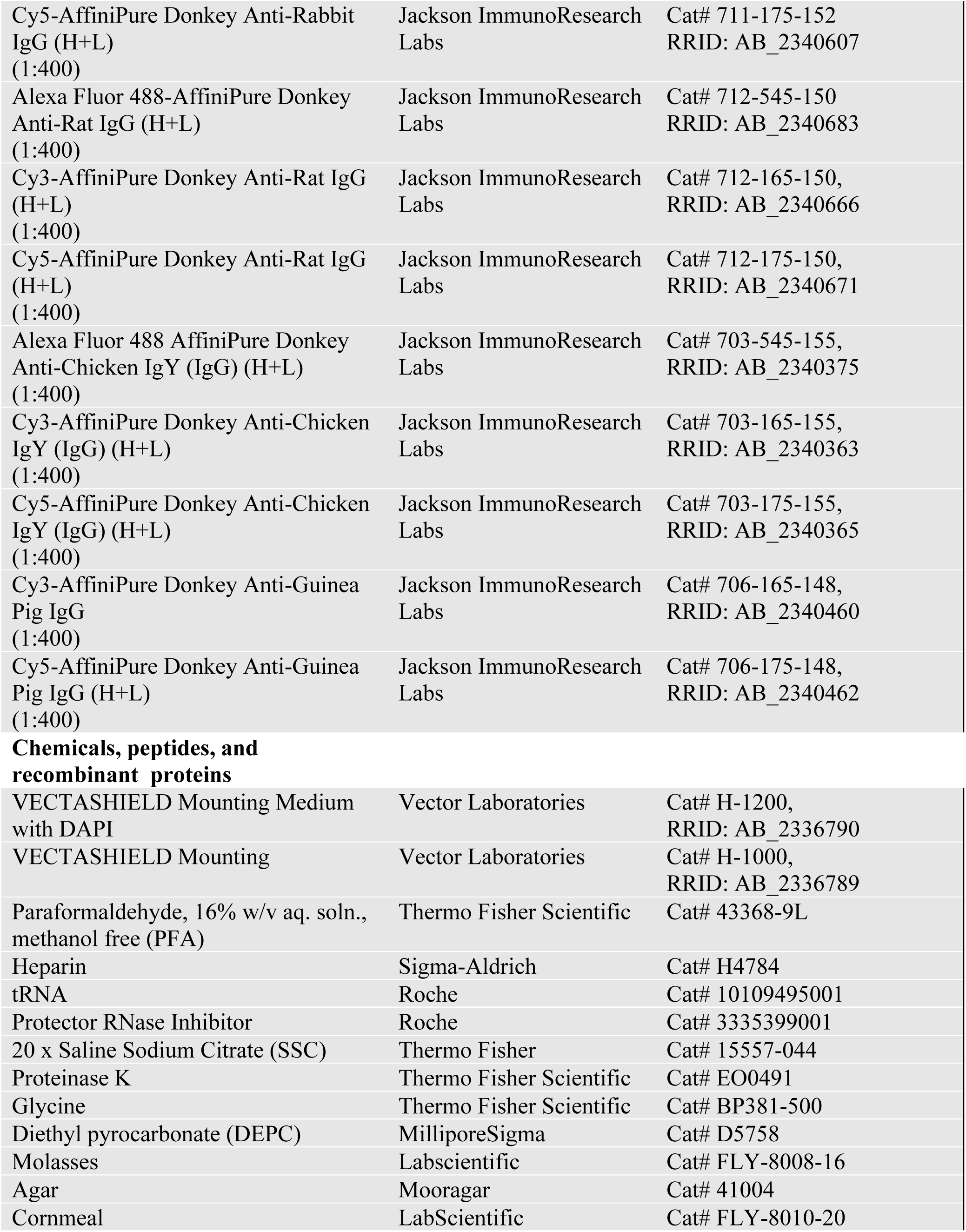
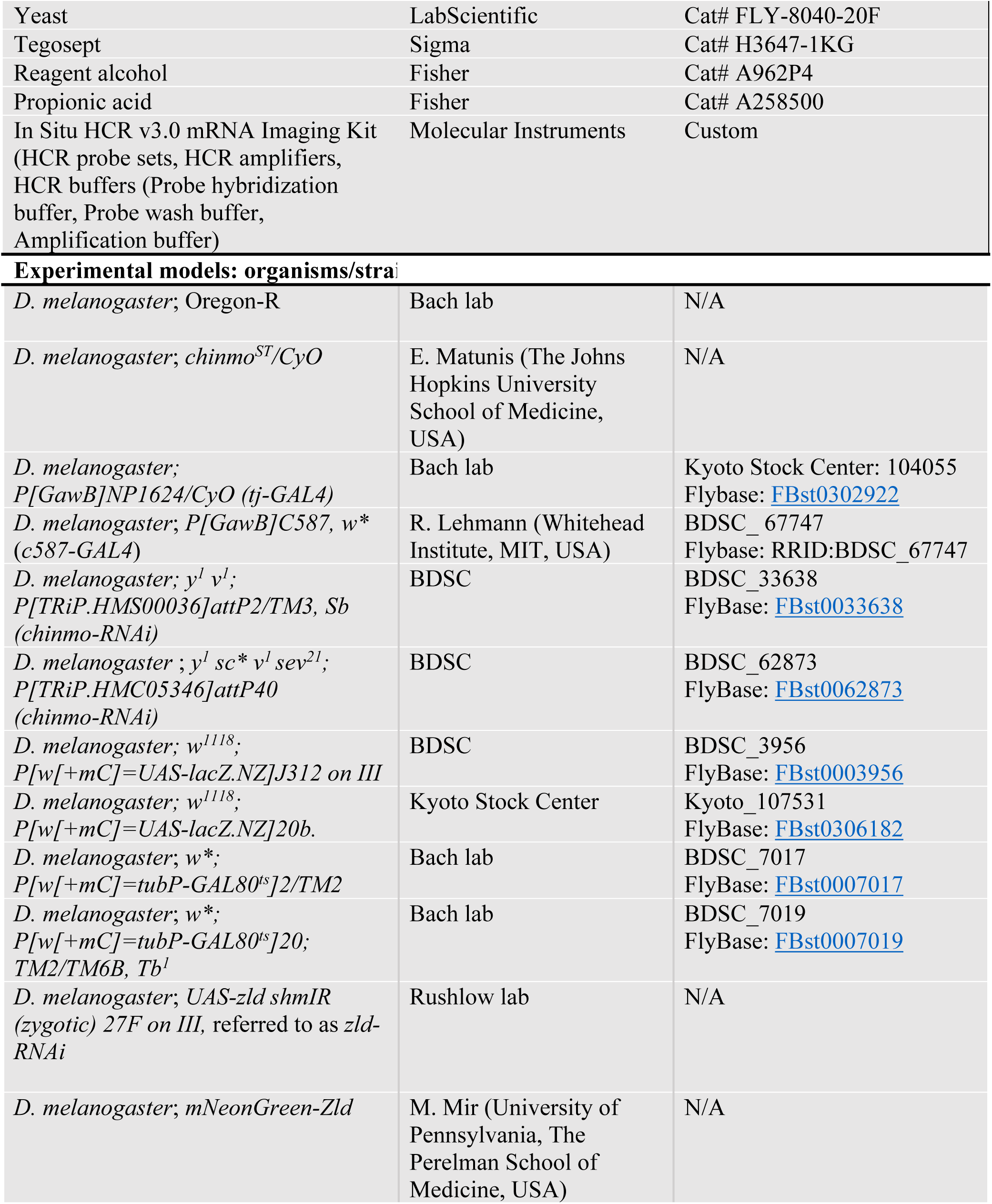
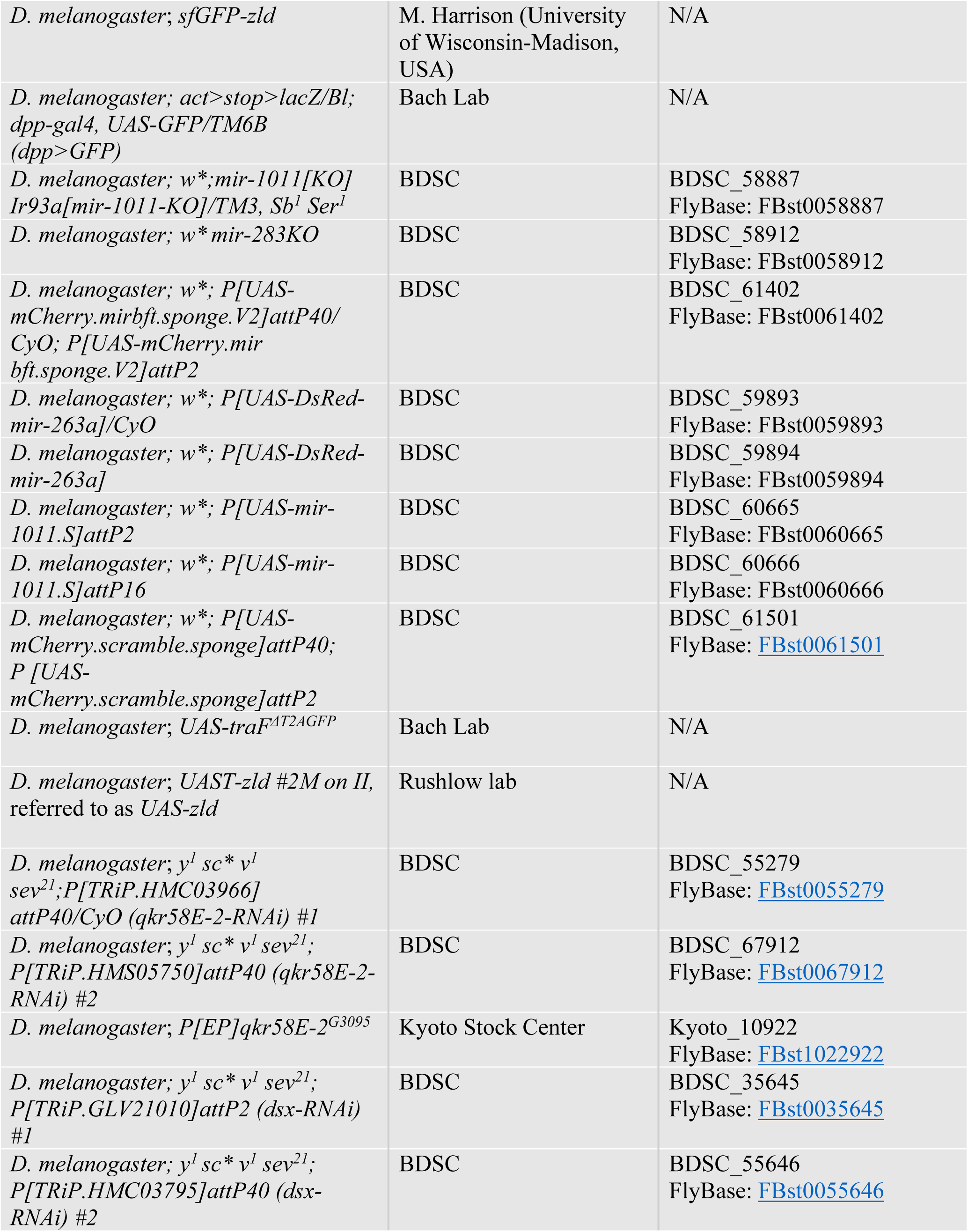
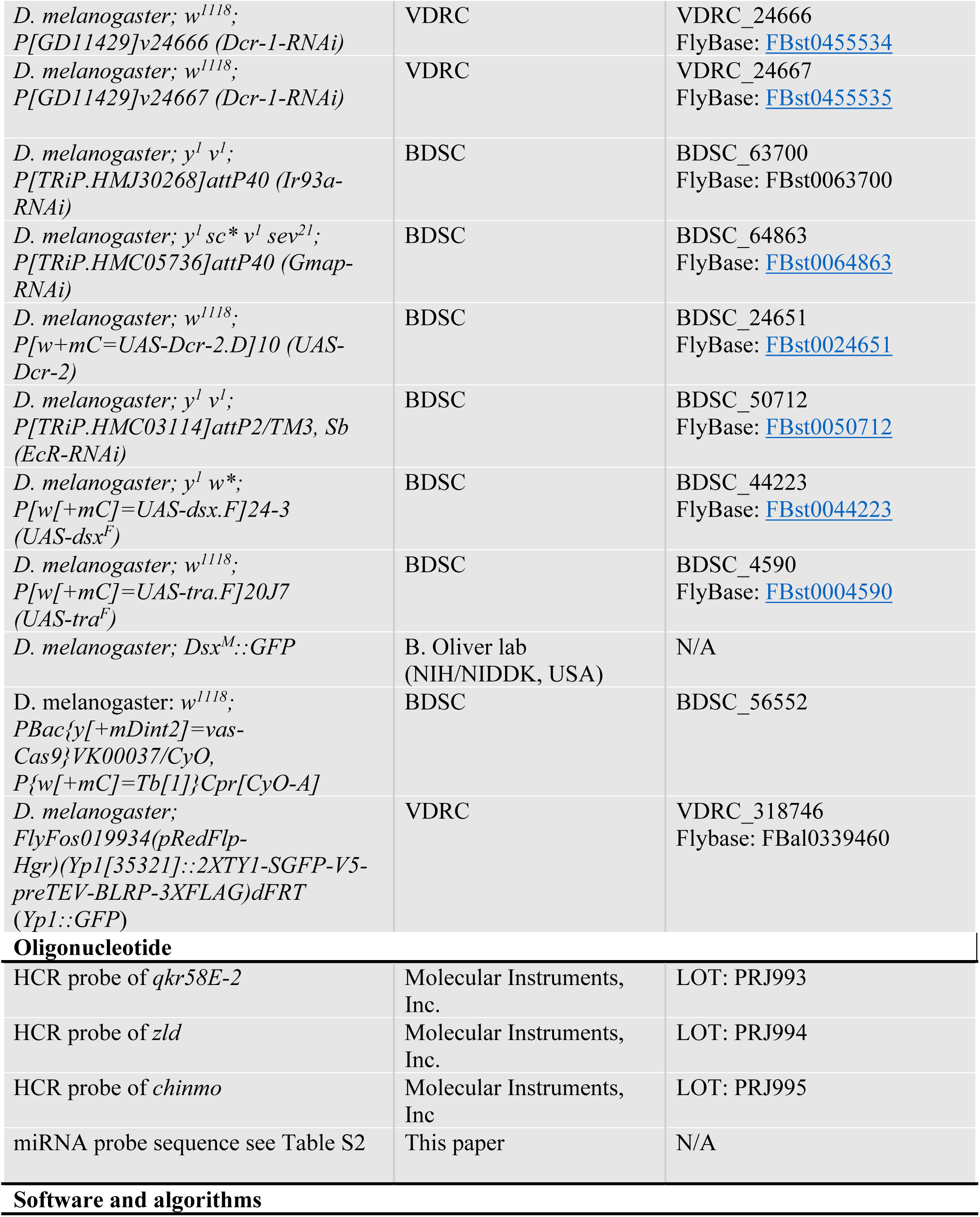
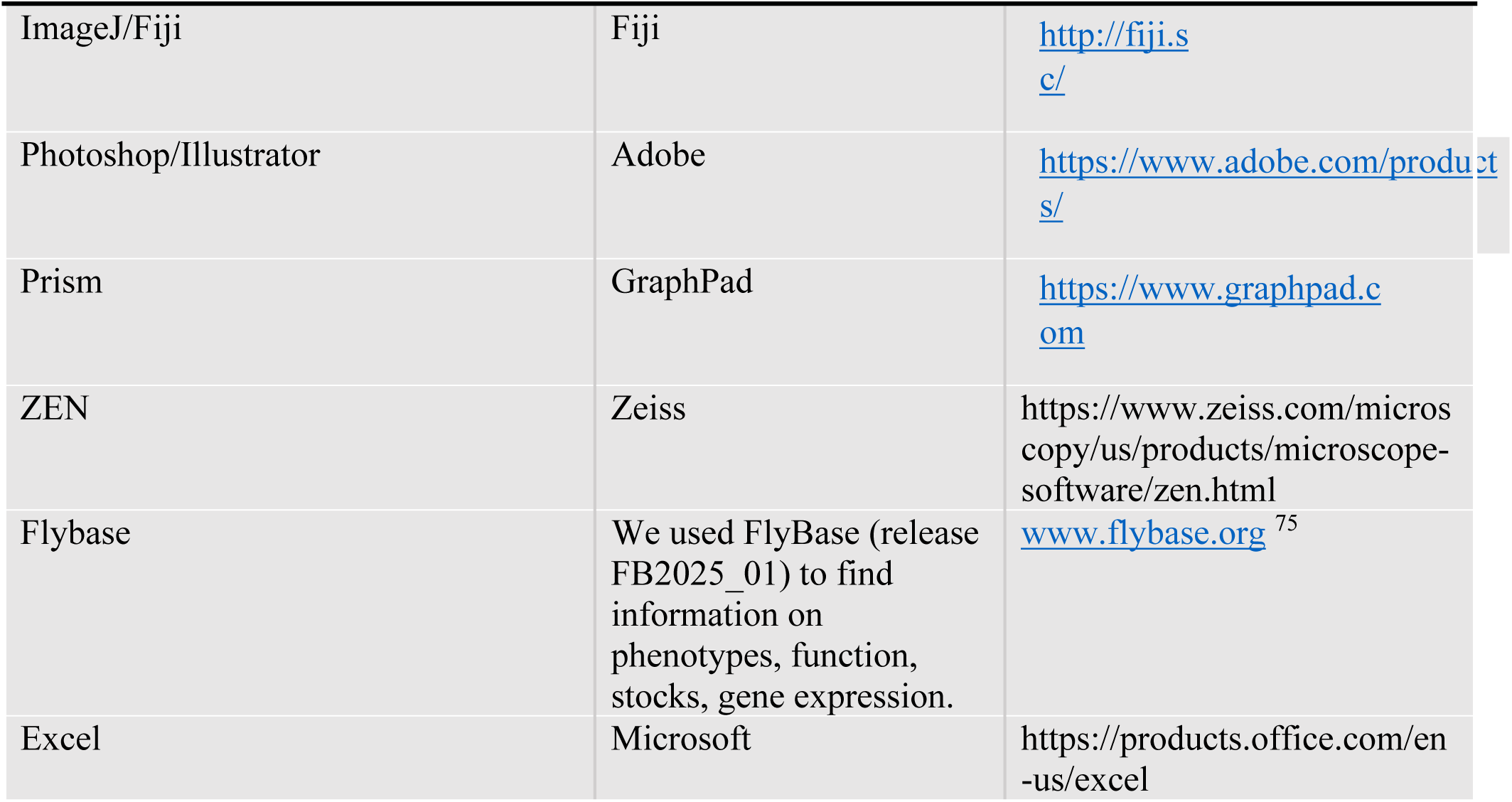

## RESOURCE AVAILABILITY

### Lead contact

Further information and requests for resources and reagents should be directed to and will be fulfilled by the Lead Contact, Erika Bach.

### Materials availability

This study did generate new unique reagents. All *Drosophila* stocks generated in this study are available from the Lead Contact without restriction.

### Data and code availability

All data reported in this paper will be shared by the Lead Contact upon request. This paper does not report original code.

Any additional information required to reanalyze the data reported in this paper is available from the Lead Contact upon request.

## EXPERIMENTAL MODEL AND SUBJECT DETAILS

### Fly lines and maintenance

*Drosophila melanogaster* strains used in this study are listed in the Key Resources Table. *Drosophila* were reared on food made with these ingredients: 1800mL molasses (LabScientific, Catalog no. FLY-8008-16), 266 g agar (Mooragar, Catalog no. 41004), 1800 g cornmeal (LabScientific, Catalog no. FLY-8010-20), 744g Yeast (LabScientific, Catalog no. FLY-8040-20F), 47 L water, 56 g Tegosept (Sigma no. H3647-1KG), 560mL reagent alcohol (Fisher no. A962P4), and 190mL propionic acid (Fisher no. A258500). Flies were raised at 25°C except crosses with *Gal80^ts^*, which were maintained at 18°C until eclosion, and the adult flies were transferred to 29°C.

We used the following *Drosophila* stocks: *tj-Gal4* (Kyoto #104055), *UAS-lacZ* (BDSC #3956), *UAS-lacZ* (Kyoto #107531), *chinmo^ST^* ^14^, *UAS-chinmo-RNAi* #1 (BDSC #33638) and #2 (BDSC #62873), *UAS-zld-RNAi* (this study), *sfGFP-zld* ^51^, *mNeonGreen-zld* ^52^, *UAS-zld* (this study), *UAS-Ir93a-RNAi* (BDSC #63700), *UAS-dicer-1-RNAi #*1 (VDRC #24666) and #2 (VDRC #24667), *miR-1011KO* (BDSC #58887), *miR-283KO* (BDSC #58912), *UAS-miR-263a-SP* (BDSC #61402), *UAS-miR-263a* #1 (BDSC #59893) and #2 (BDSC #59894), *UAS-miR-1011* #1 (BDSC #60665) and #2 (BDSC #60666); *UAS-Scramble-SP* (BDSC #61501), *UAS-traF^ΔT2AGFP^* ^15^*, tub-Gal80^ts^* ^53^, *UAS-qkr58E-2-RNAi* #1 (BDSC #55279) and #2 (BDSC #67912), *[EP]qkr58E-2^G^*^3095^ (Kyoto # 10922), *UAS-dsx-RNAi* #1 (BDSC #35645), *UAS-dsx-RNAi* #2 (BDSC 55646), Dsx^M^::GFP (this study), Yp1::GFP (VDRC#318746).

For a list of full genotypes by figure, see Table S1.

### Antibodies

To generate the Rabbit anti Zld-N, we used Pocono Rabbit Farm. Rabbits were injected with recombinant Zelda amino acids 1-618, which includes zinc fingers 1 and 2. The Zld-N antibody was pre-absorbed on *zld^M-Z-^* embryos prior to use.

### UAS-zld shmIR (zygotic) 27F transgene

The *UAS-zld shmIR (zygotic) 27F* (abbreviated *zld*-RNAi) for depleting *zld* zygotically was constructed by inserting the passenger strand sequence “CAGCAGCTACATCAACAGCTA” from ^39^ into the *Valium20* vector ^76^ and was injected into *Drosophila* embryos.

### *UAS-zld* transgene

A cDNA encoding full-length Zld was sub-cloned into *UAST* vector ^77^, which was injected into *Drosophila* embryos by Best Gene Inc.

### *Dsx^M^::FLAG-EGFP* allele (abbreviated Dsx^M^::GFP)

We designed a *dsx* locus modification to add FLAG C-terminal peptide tag (GenBank: KX714724.1) to the endogenous Dsx^M^ protein (see **Figure S10**). As a reporter of *dsx* expression, we also appended the self-cleaving T2A sequence (GenBank: MW331579.1) to the locus followed by EGFP fluorescent protein coding sequence (GenBank: MN517551.1) and a nuclear localization sequence. Because the EGFP protein is small, it can diffuse back into the cytoplasm 78. Because the Dsx proteins have a common 5’ encoded DNA-binding domain, followed by 3’ sex-specific splicing events encoding distinct effector domains, the tagged endogenous Dsx^M^ protein should be sex-specific.

The *DSX-M-FLAG-EGFP* DNA cassette was synthesized and cloned into the *pUC57-Brick* vector. We transformed flies using homology-mediated CRISPR by injecting homology DNAs into embryos of a strain expressing the Cas9 endonuclease (BDSC-56552) (*w*^1118^*; PBac{y[+mDint2]=vas-Cas9}VK00037/CyO, P{w[+mC]=Tb[1]}Cpr[CyO-A]*). gRNAs were cloned into *puc19-3xP3-dsred-attb-gypsy-U63-gRNA* and were injected into embryos. We did preliminary screening of possible new *dsx* alleles by crossing single emerging G0 flies to balancer stock (*w+; +/+; TM3,Sb/TM6B,Tb*). G1 Male flies over the balancer chromosome were crossed back to balancer stock (*w+; +/+; TM3,Sb/TM6B,Tb*). Cas9 is marked with 3xP3-GFP which is expressed in the eye, and stocks were selected not to have GFP in eye as an evidence of Cas9 removal. Multiple flies were subjected to PCR screen to identify positive insertion.

We used these primers to amplify PCR product from single fly gDNA. These primers are on either side of the tag.

For left arm: 641bp

DSXM6Fw-cgcataacttctgttaatccccagctcg

DSXMSeqRV1-accaccccggtgaacagctcctcg

Sequenced by: DSXMSeqFw1-aaatcgcactgtagcccagatctac

For right arm:378bp

EGFP-C-CATGGTCCTGCTGGAGTTCGTG

DSXM5Rv-atgtcgatctgttcctcgatttcaa

Sequenced by: DSXMSeqRV2-taaggaacgtaaggaagtgagaac

DSX-M:

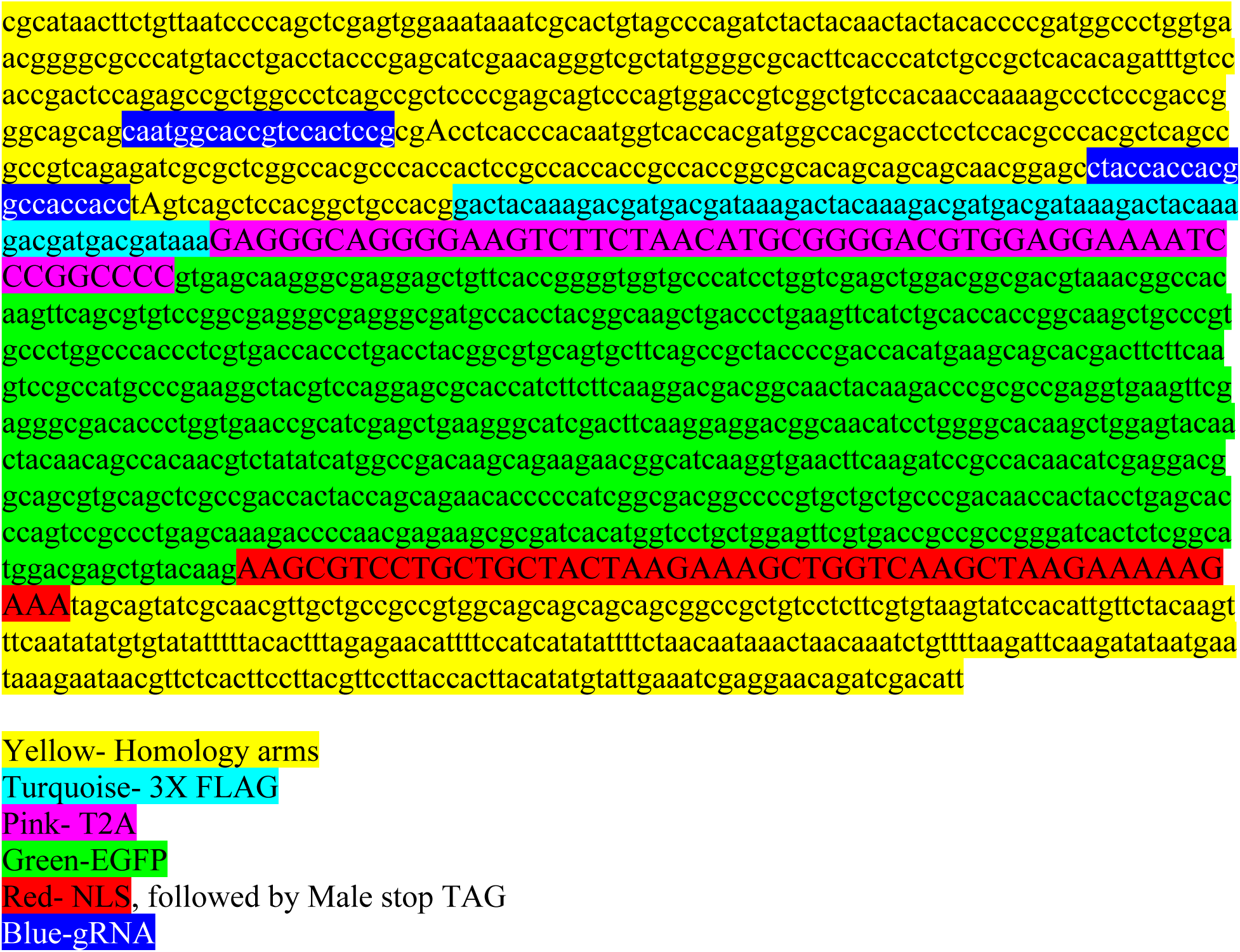

## METHODS DETAILS

### Drosophila genetics

Lineage-wide misexpression or depletion was achieved using the Gal4/UAS system ^77^. A *Gal80^ts^*transgene was used with *c587-Gal4* or *tj-Gal4* to deplete Chinmo, Qkr58E-2, Zld, EcR, or Dsx or to misexpress Zld, Qkr58E-2, Tra^F^, Dsx^F^, *miR-1011* and *miR-263a* in adult male somatic cells. *c587-Gal4* driver is active in the somatic cells of germarium including FSCs that give rise to the adult follicle epithelium ^79^.

### Dissection and immunofluorescence of adult gonads

Dissections of adult testes and ovaries were carried out as previously described ^80^. Briefly, testes/ovaries were dissected in 1X phosphate buffered saline (PBS), fixed for 30 minutes in 4% paraformaldehyde (PFA) in 1x PBS (testes). The ovaries were fixed for 30 minutes in 1x PBS with 4% PFA and 0.2% Triton X-100. The samples were washed two times for 30 minutes each at 25°C in 1x PBS with 0.5% Triton X-100, and blocked in PBTB (1x PBS, 0.2% Triton X-100 and 1% bovine serum albumin (BSA)) for 1 hour at 25°C. Primary antibodies were incubated overnight at 4°C. The samples were washed two times for 30 minutes each in 1x PBS with 0.2% Triton X-100 and incubated for 2 hours in secondary antibody in PBTB at 25°C and then washed two times for 30 minutes each in 1x PBS with 0.2% Triton X-100. They were mounted in Vectashield or Vectashield with DAPI (Vector Laboratories). Confocal images were captured using Zeiss LSM 700 and LSM 800 microscopes with a 63X objective.

For EcR detection with Ag10.2, WT ovaries, WT testes, *chinmo*-mutant testes and zld misexpressing testes were dissected on the same day, stained with the same master mix of primary and secondary antibodies, and imaged on a confocal microscope using the same settings.

### Dissection and immunofluorescence of larval tissues Wing imaginal disc and ventral nerve cord

For the dissection of larval wing imaginal discs and ventral nerve cords, properly staged larvae were rinsed in 1x PBS for immunofluorescence. The posterior part of the cuticle was carefully removed using forceps, and the specimens were inverted. The intestines and fat body were gently removed while keeping other organs intact and attached to the cuticle. Imaginal discs were dissected in 1x PBS and fixed for 30 minutes in 1x PBS containing 4% PFA and 0.2% Triton X-100 on a nutator. Following fixation, samples were washed three times for 10 minutes each in 1x PBS with 0.2% Triton X-100 and then blocked in PBTB (1x PBS with 0.2% Triton X-100 and 1% BSA) for 1 hour at room temperature. Primary antibodies were diluted in PBTB and incubated overnight at 4°C. After three 10-minute washes in 1x PBS with 0.2% Triton X-100, samples were incubated with secondary antibodies in PBTB for 2 hours at room temperature. The samples were then washed three times for 10 minutes each in 1x PBS with 0.2% Triton X-100 before mounting in Vectashield or Vectashield with DAPI (Vector Laboratories). Confocal images were acquired using Zeiss LSM 700 and LSM 800 microscopes with a 40X objective.

### Larval gonads

Male and female larvae were sexed based on gonad morphology. Male testes appeared as large, clear ovals embedded in the posterior third of the fat body, whereas female ovaries were smaller, clear, round spheres located in the same region. Larvae were dissected in 1X PBS, with the fat body and gonads carefully teased out while keeping the fat body attached to the larval cuticle. Other tissues, including the intestine, were removed.

The fat body with gonads was fixed in 4% PFA and 0.3% Triton X-100 in 1x PBS for 30 minutes with gentle rotation. Samples were washed twice for 10 minutes each in 1% PBST (1X PBS + 1% Triton X-100) and blocked in PBTB (1% PBST + 5% BSA) for 2 hours at room temperature. Primary antibodies were diluted in PBTB and incubated overnight at 4°C. Following three 20-minute washes in 0.3% PBST at room temperature, samples were incubated with secondary antibodies in PBTB for 2 hours at room temperature. After a final 20-minute wash in 0.3% PBST, samples were mounted in Vectashield or Vectashield with DAPI (Vector Laboratories). Confocal images were acquired using Zeiss LSM 700 and LSM 800 microscopes with 40X or 63X objectives.

### Hybridization Chain Reaction-Fluorescent *in situ* Hybridization (HCR-FISH)

All steps were done using RNase-free reagents and supplies with gentle rotation. The protocol for immunostaining with HCR-FISH was adapted from ^81,82^. The HCR probe set against *zld*,*qkr58E-2* and *chinmo* were purchased from Molecular Instruments, Inc. Briefly, testes/ovaries were fixed in 4% PFA in 0.1% Triton X-100 in 1xPBS-DEPC for 30 minutes at 25°C, washed with 0.5% Triton X-100 in 1x PBS-DEPC two times for 30 minutes at 25°C. Samples were blocked in 0.1% Triton X-100 in 1x PBS-DEPC with 50 µg/mL heparin and 250 µg/mL yeast tRNA (buffer hereafter called “PBTH”), and then they were then incubated with primary antibodies overnight at 4°C. The next day, the samples were washed twice in PBTH for 30 minutes. Samples were then incubated with fluorescently-labeled secondary antibodies in PBTH for 2 hours at 25°C. Samples were washed in PBTH twice for 30 minutes at 25°C. Samples were then dehydrated and rehydrated with a series of ethanol washes (25%, 50%, 75%, 100%) in 1x PBS-DEPC for 10 minutes at 25°C. Samples were treated for 7 minutes with 50 µg/mL Proteinase K, which was then inactivated by washing with 0.2% glycine twice in 1x PBS-DEPC for 5 minutes at 25°C. After Proteinase K treatment, the samples were fixed again in 4% PFA in 1x PBS-DEPC for 30 minutes at 25°C. The re-fixed samples were pre-hybridized in hybridization buffer provided by Molecular Instruments Inc. for 10 minutes at 25°C and then incubated with HCR probes overnight (12 - 16 hours) at 37°C. Samples were then washed 6 times 10 minutes at 37°C with wash buffer provided by Molecular Instruments Inc. and then twice for 5 minutes in 5x SCC at 25°C. Samples were incubated in amplification buffer provided by Molecular Instruments Inc. for 5 minutes at 25°C. The secondary reagents called “Hairpin h1 DNA” and “Hairpin h2 DNA” were prepared by heating each for 90 seconds at 95°C and cooling them at 25°C in a dark drawer for 30 minutes. Hairpin h1 DNA and Hairpin h2 DNA were mixed together at a 1:1 ratio and then added to the samples, which were then incubated in the dark environment overnight (16 hours) at 25°C. Samples were washed 6 times for 5 minutes with 5x SSC at 25°C and mounted in Vectashield plus DAPI for confocal analysis.

### Quantification of Zld protein, *zld* mRNA *and qkr58E-2* mRNA

For Zld and GFP intensity quantifications, control and experimental samples were dissected and processed in parallel on the same day to minimize variability. Images were acquired using identical microscope settings. Regions of interest (ROIs) were drawn around somatic nuclei, identified by Tj or Zfh-1 signal, and mean fluorescence intensity was measured using the freehand selection tool in ImageJ. Background signal was subtracted using a defined background ROI. Appropriate thresholding was applied to ensure accurate quantification of fluorescence intensity. Data are presented as relative fluorescence intensity, normalized to the average control values.

For *zld* and *qkr58E-2* mRNA quantification, ROIs were similarly drawn around somatic nuclei based on Traffic jam (Tj) or Zfh-1 expression, and mean fluorescence intensity for *zld* and *qkr58E-2* was measured separately in ImageJ.

### Single-molecule FISH (miRNA *in situ* hybridization)

All steps were done using RNase-free reagents and supplies with gentle rotation. The protocol for single-molecule FISH was adapted from ^83^. Testes were dissected in 1x PBS and fixed in 4% formaldehyde in 1x PBS for 20 minutes. Testes were then washed twice for 5 minutes each in 1x PBS with 0.1% Tween 20 followed by permeabilization in 1x PBS with 0.1% Triton X-100 for 2 hours. Next, the testes were washed twice with 5x SSCT (0.1% Tween 20) for 5 minutes each. Probes were added to pre-warmed Probe Hybridization Buffer (Molecular Instruments) to a final concentration of 1 µM. A total of 100 µL of hybridization solution was added to each sample, pipetted to mix, and allowed to hybridize overnight at 37°C. Samples were then washed four times with Probe Wash Buffer (Molecular Instruments) for 15 minutes each at 37°C. During the first wash, amplifier hairpins (Molecular Instruments) were individually incubated at 95°C for 2 minutes and allowed to slowly cool to room temperature. Hairpins were then added to Amplification Buffer (Molecular Instruments) to a concentration of 60 nM. After washing the samples twice with 5x SSCT for 5 minutes each, 100 µL of hairpin solution was added and the samples were allowed to incubate at room temperature overnight. The testes were then washed twice with 5x SSCT for 30 minutes each at room temperature, mounted in Vectashield with DAPI, and imaged on Zeiss LSM 800 microscope with a 63X objective and processed using ImageJ and Adobe Photoshop softwares.

### miRNA *in situ* probes and hairpin amplifiers

DNA oligonucleotides were obtained from Integrated DNA Technologies (IDT), with sequences listed in Supplementary Table S2. DNA oligonucleotide probes were designed to contain a 20– 24 nt sequence complementary to the target miRNA, flanked by initiators and linked by base linkers. Probe detection was carried out using hybridization chain reaction (HCR) with 41-nt hairpin amplifiers labeled with Cy3. Hairpin amplifiers (Set B2, conjugated to Alexa-546) were purchased from Molecular Instruments Inc.

### Quantification of farthest Zfh-1 expressing CySC

Confocal z-stacks were acquired at 1 μm intervals to encompass the entire testis. Measurements were conducted using ImageJ on a single z-section, selected based on the clear visualization of both the niche (marked by Fas3) and a CySC (Zfh-1-positive, Eya-negative cell). To determine the distance of the farthest Zfh-1-positive, Eya-negative cell, a freehand straight line was drawn from the edge of the niche to the center of the most distal Zfh-1-expressing CySC.

### Statistical Analysis

All statistical analyses were conducted in GraphPad Prism and Microsoft Excel. The specific tests, sample size, p-values and asterisks are displayed in the corresponding legends.

## Acknowledgments

We are indebted to Amelie Raz (Whitehead Institute, MIT, USA) for providing essential information about generating probes to detect miRNAs. We are grateful to Brian Oliver (NIDDK, NIH, USA) sharing unpublished reagents. Stocks obtained from the BDSC (NIH P40OD018537) were used in this study. The BDSC is supported by grant P40OD018537 from the NIH Office of Research Infrastructure Programs in collaboration with the NIGMS and NINDS. Some monoclonal antibodies was obtained from the DHSB, created by the NICHD of the NIH and maintained at The University of Iowa, Department of Biology, Iowa City, IA 52242. We thank K. White, D. Godt, P. Rangan, N. Sokol for antibodies, and M. Mir, M. Harrison for fly stocks. Work in the Bach lab was funded by R01-GM085075 from NIGMS.

## Author contributions

S.H. conducted the experiments; S.H. and E.A.B. designed the experiments and wrote the paper; H-Y.L., C.R. and P.K.B. provided critical reagents.

## Declaration of interests

The authors declare no competing interests.

## Notes

### Competing Interest Statement

The authors have declared no competing interest.

