## Supplementary Material for "The pioneer factor Zelda induces male-to-female somatic sex reversal in adult tissues"

| **Table S1: Genotypes analyzed in main body and supplementary data, related to Figs. 1-7** | | |
|---|---|---|
| **Figures** | **Full genotype** | **Abbreviation used in Figure** |
| 1A | c587-Gal4/Y; tub-Gal80$^{ts}$/+; UAS-lacZ/+ | c587$^{ts}$>lacZ |
| 1B | c587-Gal4/Y; tub-Gal80$^{ts}$/+; +/UAS-chinmo-RNAi | c587$^{ts}$>chinmo-i |
| 1C | *Oregon-R* | WT |
| 1D | w/Y; chinmo$^{ST}$/chinmo$^{ST}$ | chinmo$^{ST/ST}$ |
| 1E | c587-Gal4/Y; tub-Gal80$^{ts}$/+; UAS-lacZ/+ | c587$^{ts}$>lacZ |
| 1E | c587-Gal4/Y; tub-Gal80$^{ts}$/+; +/UAS-chinmo-RNAi | c587$^{ts}$>chinmo-i |
| 1F | *Oregon-R* | WT |
| 1F | w/Y; chinmo$^{ST}$/chinmo$^{ST}$ | chinmo$^{ST/ST}$ |
| 1G | *w/Y; tj-Gal4/UAS-lacZ; UAS-Dcr-2/UAS-lacZ* | *tj>lacZ; lacZ* |
| 1H | *w/Y; tj-Gal4/UAS-chinmo-RNAi; UAS-Dcr-2/UAS-lacZ* | *tj>chinmo-i; lacZ* |
| 1I | *w/Y; tj-Gal4/UAS-chinmo-RNAi; UAS-Dcr-2/UAS-zld-RNAi* | *tj>chinmo-i; zld-i* |
| 1J (left side) | *w/Y; tj-Gal4/UAS-lacZ; UAS-Dcr-2/UAS-lacZ* | *tj>lacZ; lacZ* |
| 1J (left side) | *w/Y; tj-Gal4/UAS-chinmo-RNAi; UAS-Dcr-2/UAS-lacZ* | *tj>chinmo-i; lacZ* |
| 1J (left side) | *w/Y; tj-Gal4/UAS-chinmo-RNAi; UAS-Dcr-2/UAS-zld-RNAi* | *tj>chinmo-i; zld-i* |
| 1J (right side) | c587-Gal4/Y; chinmo$^{ST}$/CyO | c587>chinmo$^{ST}$/CyO |
| 1J (right side) | c587-Gal4/Y; chinmo$^{ST}$/chinmo$^{ST}$ | c587>chinmo$^{ST/ST}$ |
| 1J (right side) | c587-Gal4/Y; chinmo$^{ST}$/chinmo$^{ST}$; UAS-zld-RNAi/+ | c587>chinmo$^{ST/ST}$; zld-i |
| 2A, C | *Oregon-R* | WT |
| 2B, C | w/Y; chinmo$^{ST}$/chinmo$^{ST}$ | chinmo$^{ST/ST}$ |
| 2E, F, G, N, O, P, Q | *Oregon-R* | WT |
| 2H, I, J, N, O, P | c587-Gal4/Y; tub-Gal80$^{ts}$/+; UAS-chinmo-RNAi/+ | c587$^{ts}$>chinmo-i |
| 2K, N | *w/Y; +/+; miR-1011KO/miR-1011KO* | miR mutant or miR Sponge |
| 2L, O | *w/Y; tj-Gal4/UAS-miR-263aSP; +/UAS-miR-263a-SP* | miR mutant or miR Sponge |
| 2M, P | *w, miR-283KO/Y* | miR mutant or miR Sponge |
| 2R | *w, miR-283KO/Y* | *miR-283KO* |

| | | |
|---|---|---|
| 2S | *w/Y; +/+; miR-1011KO/miR-1011KO* | *miR-1011KO* |
| 2T | *w/Y; tj-Gal4/UAS-miR-263aSP; +/UAS-miR-263a-SP* | *tj>miR-263aSP* |
| 2U | *Oregon-R* | WT |
| 2U | *w, miR-283KO/Y* | *miR-283KO* |
| 2U | *w/Y; +/+; miR-1011KO/miR-1011KO* | *miR-1011KO* |
| 2V | *w/Y; tj-Gal4/UAS-miR-Scramble-SP; +/UAS-Scramble-SP* | *tj>Scramble-SP* |
| 2V | *w/Y; tj-Gal4/UAS-miR-263aSP; +/UAS-miR-263a-SP* | *tj>miR-263aSP* |
| 3A, E, F | *Oregon-R* | WT |
| 3B, E, F | *w, miR-283KO/Y* | *miR-283KO* |
| 3C, E, F | *w/Y; tj-Gal4/UAS-miR-263aSP; +/UAS-miR-263a-SP* | *tj>miR-263aSP* |
| 3D, E, F | *w/Y; +/+; miR-1011KO/miR-1011KO* | *miR-1011KO* |
| 3G, J | *c587-Gal4/Y; tub-Gal80*$^{ts}$*/+; +/+* | *c587*$^{ts}$*>+* |
| 3H, J | *c587-Gal4/Y; tub-Gal80*$^{ts}$*/+; UAS-chinmo-RNAi/+* | *c587*$^{ts}$*>chinmo-i* |
| 3I, J | *c587-Gal4/Y; tub-Gal80*$^{ts}$*/UAS-miR-263a; UAS-miR-1011/UAS-chinmo-RNAi* | *c587*$^{ts}$*>miR-263a; miR-1011; chinmo-i* |
| 3J | *c587-Gal4/Y; tub-Gal80*$^{ts}$*/UAS-miR-263a; UAS-chinmo-RNAi/+* | *c587*$^{ts}$*>miR-263a; chinmo-i* |
| 3J | *c587-Gal4/Y; tub-Gal80*$^{ts}$*/UAS-miR-1011; UAS-chinmo-RNAi/+* | *c587*$^{ts}$*>miR-1011; chinmo-i* |
| 4B, C, F, G, I, J | *c587-Gal4/Y; tub-Gal80*$^{ts}$*/UAS-zld; UAS-traF*$^{\Delta T2AGFP}$*/+* | *c587*$^{ts}$*>traF*$^{\Delta T2AGFP}$*; zld* |
| 4D, E | *c587-Gal4/Y; tub-Gal80*$^{ts}$*/+; UAS-traF*$^{\Delta T2AGFP}$*/UAS-chinmo-RNAi* | *c587*$^{ts}$*>traF*$^{\Delta T2AGFP}$*; chinmo-i* |
| 4H, K, L, M, N | *c587-Gal4/Y; tub-Gal80*$^{ts}$*/UAS-zld; +/+* | *c587*$^{ts}$*>zld* |
| 5A-C | *c587-Gal4/Y; tub-Gal80*$^{ts}$*/+; UAS-chinmo-RNAi/+* | *c587*$^{ts}$*>chinmo-i* |
| 5D, H | *Oregon-R* | WT testis |
| 5E, H | *Oregon-R* | WT ovary |
| 5F-H | *c587-Gal4/Y; tub-Gal80*$^{ts}$*/UAS-zld; +/+* | *c587*$^{ts}$*>zld* |
| 5I, J, L | *c587-Gal4/Y; tub-Gal80*$^{ts}$*/UAS-zld; UAS-traF*$^{\Delta T2AGFP}$*/+* | *c587*$^{ts}$*>traF*$^{\Delta T2AGFP}$*; zld* |
| 5K, L | *c587-Gal4/Y; UAS-qkr58E-2-RNAi/UAS-zld; tubGal80*$^{ts}$*/UAS-traF*$^{\Delta T2AGFP}$ | *c587*$^{ts}$*>traF*$^{\Delta T2AGFP}$*; zld; qkr58E-2-i #1* |
| 5M | *c587-Gal4/Y; tub-Gal80*$^{ts}$*/+; +/+* | *c587*$^{ts}$*>+* |

| | | |
|---|---|---|
| 5M | *c587-Gal4/Y; tub-Gal80*$^{ts}$*/+; UAS-chinmo-RNAi/+* | *c587*$^{ts}$*>chinmo-i* |
| 5M | *c587-Gal4/Y; tub-Gal80*$^{ts}$*/UAS-qkr58E-2-RNAi#1; UAS-chinmo-RNAi/+* | *c587*$^{ts}$*>qkr58E-2-i #1; chinmo-i* |
| 5M | *c587-Gal4/Y; tub-Gal80*$^{ts}$*/UAS-qkr58E-2-RNAi#2; UAS-chinmo-RNAi/+* | *c587*$^{ts}$*>qkr58E-2-i #2; chinmo-i* |
| 5N, Q | *c587-Gal4/Y; +/+; UAS-traF*$^{\Delta T2AGFP}$*/+* | *c587>traF*$^{\Delta T2AGFP}$ |
| 5O-Q | *c587-Gal4/Y; +/UAS-qkr58E-2*$^{G3095}$*; UAS-traF*$^{\Delta T2AGFP}$*/+* | *c587>traF*$^{\Delta T2AGFP}$*; qkr58E-2* |
| 5R | *c587-Gal4/Y; +/+* | *c587>+* |
| 5R | *c587-Gal4/Y; +/UAS-qkr58E-2*$^{G3095}$ | *c587>qkr58E-2* |
| 6A, E | *c587-Gal4/w; tub-Gal80*$^{ts}$*/+* | *c587*$^{ts}$*>+* ovary |
| 6B, E | *c587-Gal4/w; tub-Gal80*$^{ts}$*; EcR-i/+* | *c587*$^{ts}$*>EcR-i* ovary |
| 6C, E | *c587-Gal4/Y; tub-Gal80*$^{ts}$*/+* | *c587*$^{ts}$*>+* testis |
| 6D, E | *c587-Gal4/Y; tub-Gal80*$^{ts}$*/+; UAS-chinmo-RNAi/+* | *c587*$^{ts}$*>chinmo-i* testis |
| 6F-I | *c587-Gal4/Y; tub-Gal80*$^{ts}$*/UAS-zld; +/+* | *c587*$^{ts}$*>zld* |
| 7A, B | Dsx$^{M}$::GFP/*Tb* | |
| 7C, D | Yp1::GFP/*CyO* | |
| 7E, F, H, I, J | *c587-Gal4/Y; tub-Gal80*$^{ts}$*/+; UAS-traF*$^{\Delta T2AGFP}$*/+* | *c587*$^{ts}$ *>traF*$^{\Delta T2AGFP}$ |
| 7G, H, I, J | *c587-Gal4/Y; tub-Gal80*$^{ts}$*/UAS-zld; UAS-traF*$^{\Delta T2AGFP}$*/+* | *c587*$^{ts}$*>traF*$^{\Delta T2AGFP}$*; zld* |
| **Supplementary Figures** | | |
| S2A | *w/Y; +/+; dpp-Gal4, UAS-GFP/+* | *dpp>+* |
| S2B | *w/Y; +/+; dpp-Gal4, UAS-GFP/UAS-zld-RNAi* | *dpp>zld-i* |
| S2C, D, G, H, I | *w/Y; tj-Gal4/+; +/+* | *tj>+* |
| S2E, G, I | *w/Y; tj-Gal4/+; +/UAS-zld-RNAi* | *tj>zld-i* |
| S2F, H | *w/Y; tj-Gal4/+; UAS-zld/+* | *tj>zld* |
| S2J-L | sfGFP-Zld/Y | |
| S2N-P | mNeonGreen-Zld/Y | |
| S2M | sfGFP-Zld/Y; *chinmo*$^{ST}$*/+* | *sfGFP-Zld/Y*; *chinmo*$^{ST}$*/+* |
| S2M | *FM7/Y*; *chinmo*$^{ST}$*/chinmo*$^{ST}$ | *FM7/Y*; *chinmo*$^{ST}$*/chinmo*$^{ST}$ |
| S2M | sfGFP-Zld/Y; *chinmo*$^{ST}$*/chinmo*$^{ST}$ | *sfGFP-Zld/Y*; *chinmo*$^{ST}$*/chinmo*$^{ST}$ |
| S2Q | *FM7/Y; tj-Gal4/+; +/+* | *FM7/Y; tj>+* |
| S2Q | *FM7/Y; tj-Gal4/+; UAS-dcr-2/UAS-chinmo-RNAi* | *FM7/Y; tj>chinmo-i* |

| | | |
|---|---|---|
| S2Q | mNeonGreen-Zld*/Y; tj-Gal4/+; UAS-dcr-2/UAS-chinmo-RNAi* | *mNG-Zld/Y; tj>chinmo-i* |
| S3A | *Oregon-R* | WT adult ovary |
| S3B | *Oregon-R* | WT larval ovary |
| S3C | *Oregon-R* | WT larval testis |
| S3D | *Oregon-R* | WT larval VNC |
| S3E | *c587-Gal4/w; tub-Gal80*$^{ts}$*/+; +/UAS-zld-RNAi* | *c587*$^{ts}$*>zld-i* |
| S4A, C | *w/Y; tj-Gal4/+; +/+* | *tj>+* |
| S4B, C | *w/Y; tj-Gal4/+; +/UAS-Dicer-1-RNAi* | *tj>Dcr-1-i* |
| S5A, C | *w/Y; tj-Gal4/+; +/+* | *tj>+* |
| S5B, C | *w/Y; tj-Gal4/+; UAS-Ir93a-RNAi* | *tj>Ir93a-i* |
| S6A, C | *w/w; tj-Gal4/+; +/+* | *tj>+* |
| S6B, C | *w/w; tj-Gal4/UAS-qkr58E-2-RNAi #1; +/+* | *tj>qkr58E-2-i #1* |
| S6C | *w/w; tj-Gal4/UAS-qkr58E-2-RNAi #2; +/+* | *tj>qkr58E-2-i #2* |
| S6D, F | *c587-Gal4/Y; +/+; +/+* | *c587>+* |
| S6E, F | *c587-Gal4/Y; +/UAS-qkr58E-2*$^{G3095}$ | *c587>qkr58E-2*$^{G3095}$ |
| S7A, C | *w/w; tj-Gal4/+; UAS-traF*$^{\Delta T2AGFP}$*/+* | *tj>traF*$^{\Delta T2AGFP}$ |
| S7B | *w/w; tj-Gal4/UAS-qkr58E-2-RNAi; UAS-traF*$^{\Delta T2AGFP}$*/+* | *tj>traF*$^{\Delta T2AGFP}$*; qkr58E-2-i #1* |
| S7C | *w/w; tj-Gal4/UAS-qkr58E-2-RNAi #1; UAS-traF*$^{\Delta T2AGFP}$*/+* | *tj>traF*$^{\Delta T2AGFP}$*; qkr58E-2-i #1* |
| S7C | *w/w; tj-Gal4/UAS-qkr58E-2-RNAi #2; UAS-traF*$^{\Delta T2AGFP}$*/+* | *tj>traF*$^{\Delta T2AGFP}$*; qkr58E-2-i #2* |
| S7C | *c587-Gal4/w; +/+; UAS-traF*$^{\Delta T2AGFP}$*/+* | *c587>traF*$^{\Delta T2AGFP}$ |
| S7C | *c587-Gal4/w; +/UAS-qkr58E-2-RNAi #1; UAS-traF*$^{\Delta T2AGFP}$*/+* | *c587>traF*$^{\Delta T2AGFP}$*; qkr58E-2-i #1* |
| S7C | *c587-Gal4/w; +/UAS-qkr58E-2-RNAi #2; UAS-traF*$^{\Delta T2AGFP}$*/+* | *c587>traF*$^{\Delta T2AGFP}$*; qkr58E-2-i #2* |
| S7D, G | *w/w; tj-Gal4/+; +/+* | *tj>+* |
| S7E | *w/w; tj-Gal4/UAS-qkr58E-2-RNAi; +/+* | *tj>qkr58E-2-i* |
| S7F | *w/w; tj-Gal4, tub-Gal80*$^{ts}$*/UAS-dsx-RNAi; UAS-Dcr-2/+* | *tj*$^{ts}$*>dsx-i* |
| S7G | *w/w; tj-Gal4/UAS-qkr58E-2-RNAi#1; +/+* | *tj>qkr58E-2-i #1* |
| S7G | *w/w; tj-Gal4/UAS-qkr58E-2-RNAi#2; +/+* | *tj>qkr58E-2-i #2* |
| S7G | *w/w; tj-Gal4, tub-Gal80*$^{ts}$*/UAS-dsx-RNAi#1; UAS-Dcr-2/+* | *tj*$^{ts}$*>dsx-i #1* |

| | | |
|---|---|---|
| S7G | *w/w; tj-Gal4, tub-Gal80$^{ts}$/UAS-dsx-RNAi#2; UAS-Dcr-2/+* | *tj$^{ts}$>dsx-i #2* |
| S8A, C | *w/Y; tj-Gal4/+; +/+* | *tj>+* |
| S8B, C | *w/Y; tj-Gal4/UAS-qkr58E-2-i; +/+* | *tj>qkr58E-2-i #1* |
| S9A, D | *c587-Gal4/Y; tub-Gal80$^{ts}$/+* | *c587$^{ts}$>+* |
| S9B, D | *c587-Gal4/Y; tub-Gal80$^{ts}$/+; +/UAS-tra$^{F}$* | *c587$^{ts}$>tra$^{F}$* |
| S9C, D | *c587-Gal4/Y; tub-Gal80$^{ts}$/+; +/UAS-dsx$^{F}$* | *c587$^{ts}$>dsx$^{F}$* |

| | | | |
|---|---|---|---|
| **Table S2** | | | |
| **miRNA probe sequences for HCR** | | | |
| **miRNA target** | **Amplifier Set** | **Length** | **DNA oligonucleotide sequences** |
| **miRNA** | | | Initiator Probe |
| miR 1011 | 1 | 64 | CCTCGTAAATCCTCATCAATCATCCAGTAAACCGCCatataCTGCGAGCGATTTGAACCAATAA |
| miR 263a | 1 | 65 | CCTCGTAAATCCTCATCAATCATCCAGTAAACCGCCatataCCCGTGAATTCTTCCAGTGCCATT |
| miR 283 | 1 | 63 | CCTCGTAAATCCTCATCAATCATCCAGTAAACCGCCatataCCAGAATTACCAGCTGATATTT |

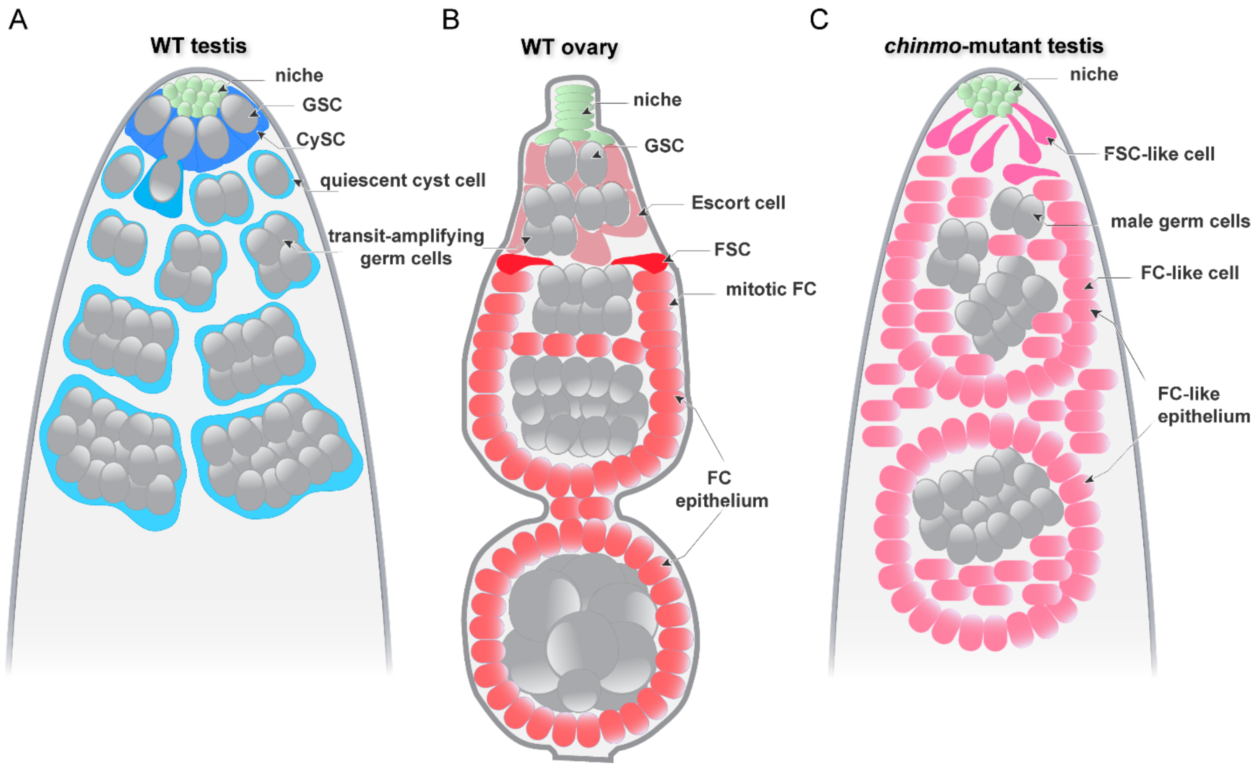

**Figure S1: Schematic of WT testis, WT ovary and feminized testis, related to Fig. 1**

(A) Cyst stem cells of a wild-type (WT) testis (CySCs, dark blue) are mitotic and reside in the same niche (green) as germline stem cells (GSCs, gray). Their daughter cells (cyst cells, light blue) are squamous, quiescent, and encyst the differentiating male germ cells (gray).

(B) Follicle stem cells (FSC, dark red) reside in the germaria of a WT ovary. They give rise to follicle cells (FC, light red) that are mitotic and epithelial. FCs form an epithelium that encyst differentiating female germ cells (gray). The ovary also has a niche (green) that supports GSCs (gray). Escort cells (ECs, pink) envelope early female germ cells.

(C) In a testis somatically depleted of *chinmo*, the niche (green) remains and the CySCs transdifferentiate into FSC-like cells (dark pink). These FSC-like cells give rise to FC-like cells (pink) that cannot properly support male germ cells, leading to defective spermatogenesis and infertility.

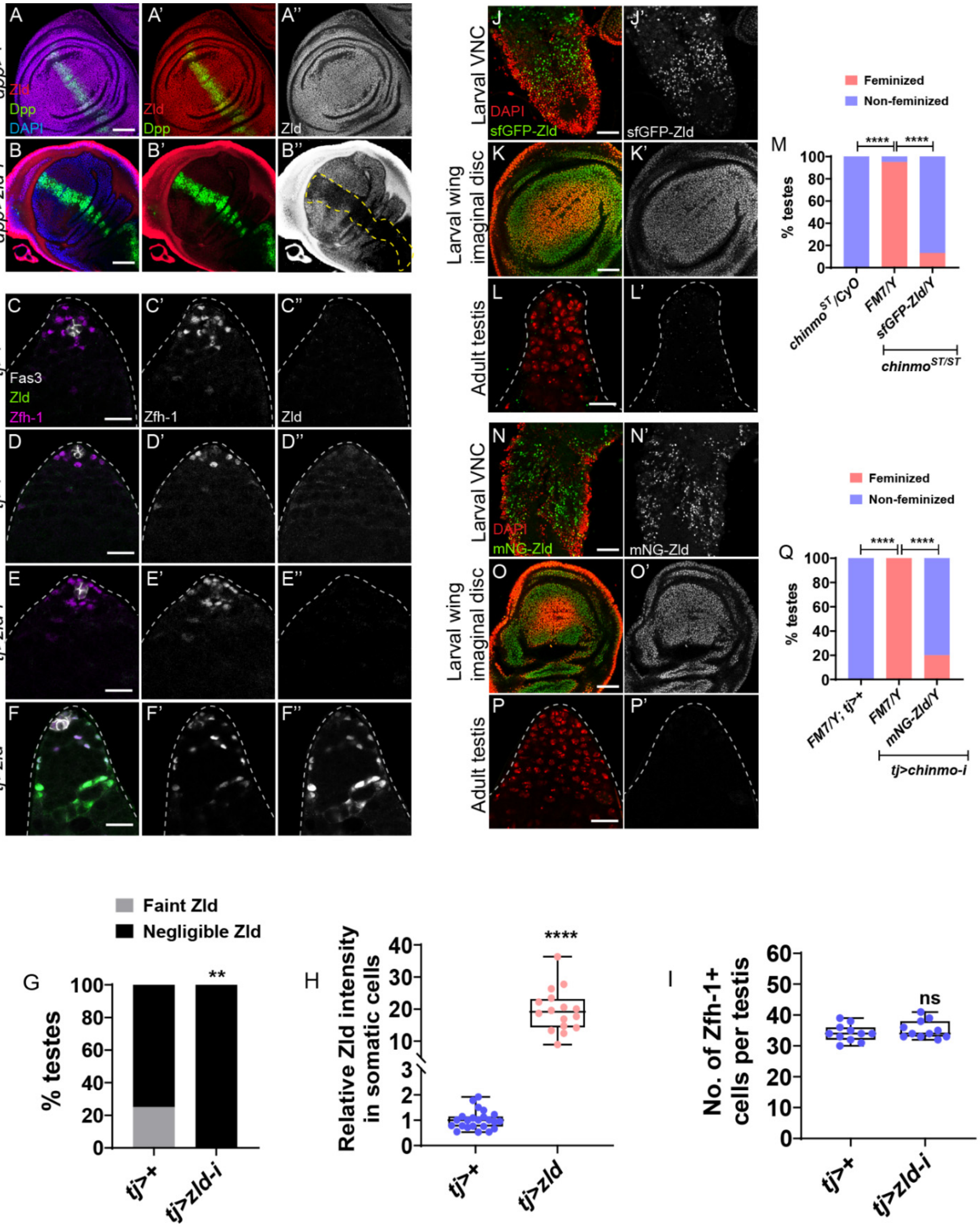

**Figure S2: Validation of Zld antibody, Zld-tagged alleles and *zld*-i line, related to Fig. 1.**

(A, B) Representative confocal images of control (*dpp>+*) (A) and *dpp>zld-i* (B) wing imaginal discs stained for Zld (red, grayscale), Dpp domain (green) and DAPI (blue). Yellow dotted lines (B'') show the diminished expression of Zld in *dpp>zld-i*.

(C-F) Representative confocal images of control (*tj*>+) displaying negligible Zld expression (C), faint Zld expression (D), *tj>zld-i* (E), and *tj>zld* (F) testes. Testes are stained for Fas3 (grayscale), Zfh-1 (magenta, grayscale) and Zld (green, grayscale).

(G) Graph showing the percentage of testes displaying faint and negligible Zld expression in *tj*>+ (n=36) and *tj>zld-i* (n=25).

(H) Graph showing relative Zld expression in the somatic cells of *tj*>+ (n=12) and *tj>zld* (n=12), testes.

(I) Graph depicting the total number of Zfh-1 positive cells in *tj*>+ (n=11) and *tj>zld-i* (n=11) testes.

(J-L) Representative confocal images of sfGFP-Zld in larval ventral nerve cord (VNC) (J), wing imaginal disc (K) and adult testis (L). sfGFP-Zld is shown in green and grayscale. DAPI is marked in red.

(M) Graph showing the percentage of feminized and non-feminized testes in *FM7/Y; chinmo*$^{ST/}$*CyO* (n=15), *FM7/Y; chinmo*$^{ST/ST}$ (n=20) and *sfGFP-Zld/Y; chinmo*$^{ST/ST}$ (n=15).

(N-P) Representative confocal images of mNG-Zld in larval VNC (N), wing imaginal disc (O) and adult testis (P). mNG-Zld is shown in green and grayscale. DAPI is marked in red.

(Q) Graph showing the percentage of feminized and non-feminized testes in *FM7/Y; tj*>+ (n=15), *FM7/Y; tj>chinmo-i* (n=20) and *mNG-Zld/Y; tj>chinmo-i* (n=20).

Statistical analysis was performed using Student's t-test (H, I) Fisher's exact test (G, M, Q) (n.s= not significant, $P > 0.05$; $^{**}P < 0.01$; $^{****}P < 0.0001$).

Scale bars: 20 μm and 50μm in J, K, N, O.

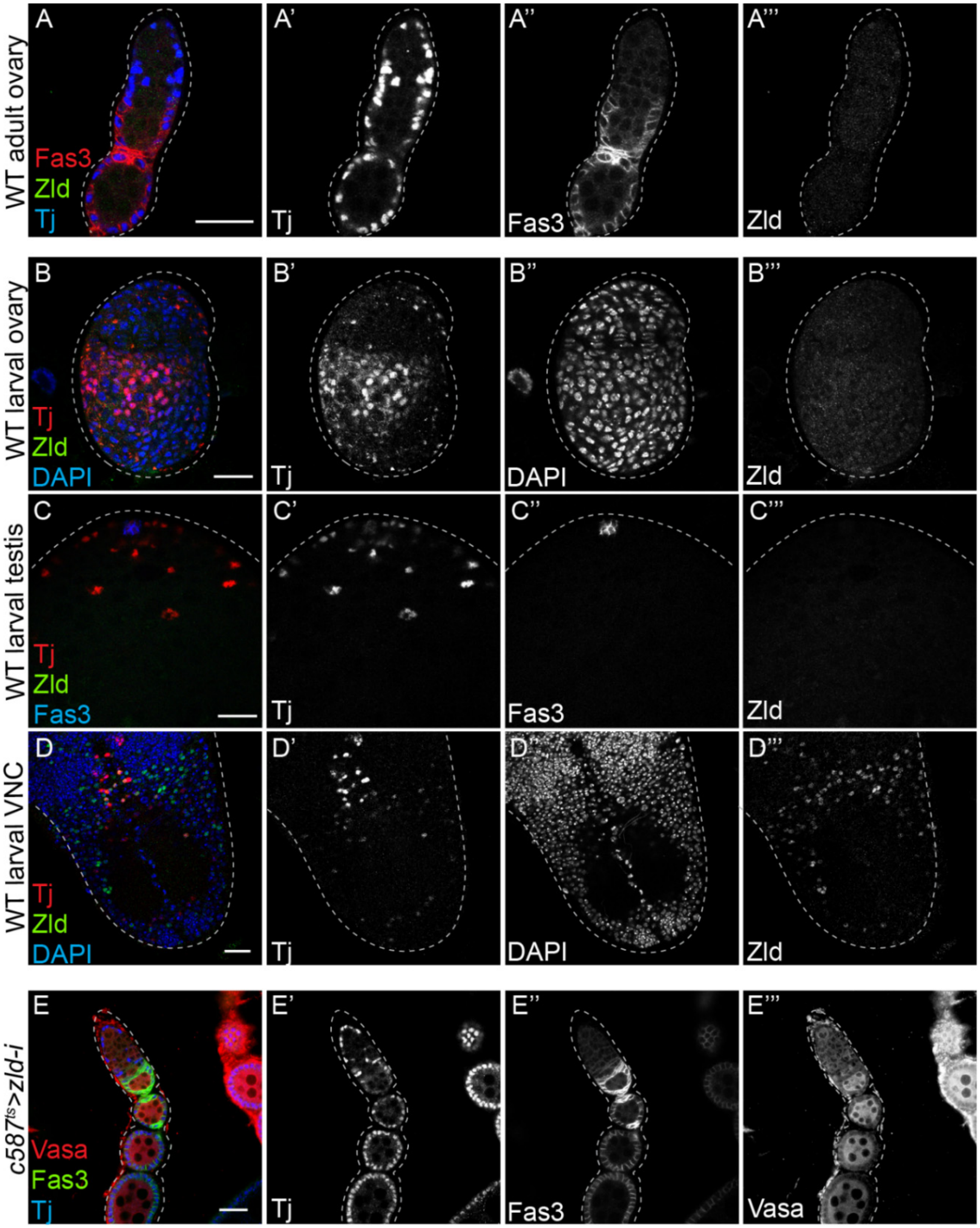

**Figure S3: Zld is not expressed in adult and larval ovary and is not required in oogenesis, related to Fig. 1.**

(A) Representative confocal images of WT adult ovariole stained for Tj (blue, grayscale), Fas3 (red, grayscale) and Zld (green, grayscale).

(B-D) Representative confocal images of WT larval ovary (B), WT larval testis (C), and WT larval VNC (D) stained for Tj (red, grayscale), DAPI (blue, grayscale), Zld (green, grayscale) and Fas3 (blue, grayscale).

(E) Representative confocal images of *c587*$^{ts}$*>zld-i* ovariole stained for Tj (blue, grayscale), Fas3 (green, grayscale) and Vasa (red, grayscale).

Scale bars: 20 µm.

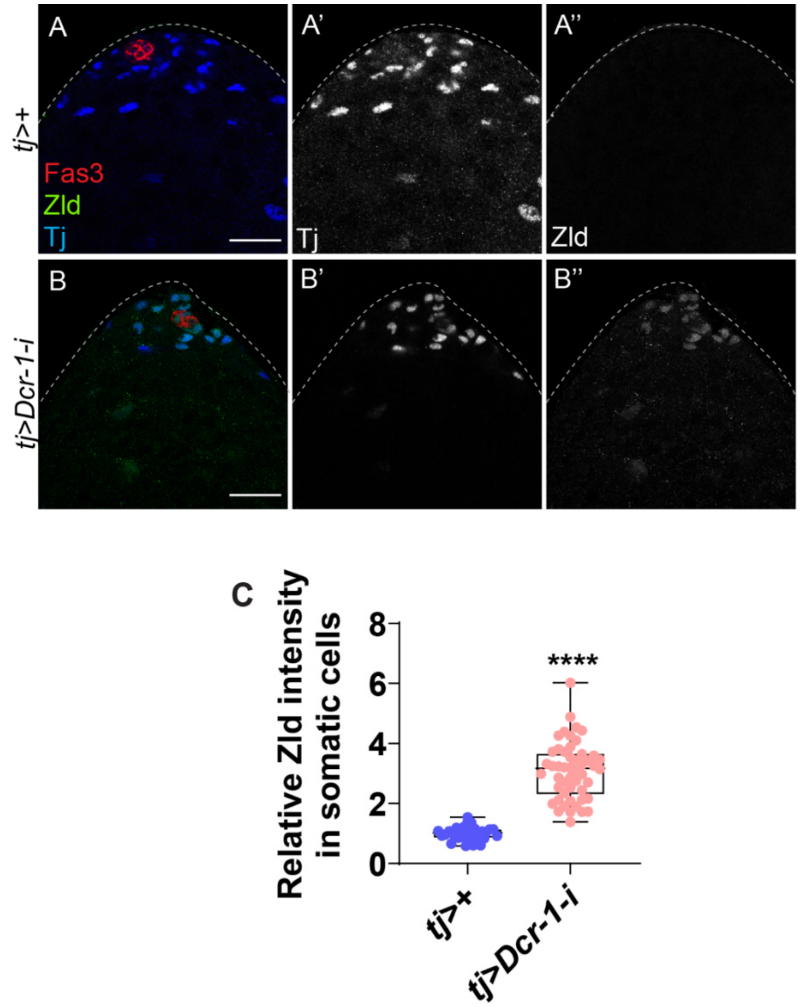

**Figure S4: Zld is upregulated in *Dcr-1*-depleted somatic cells, related to Fig. 2.**

(A, B) Representative confocal images of *tj*>+ (A), and *tj>Dcr-1-i* (B) testes stained for Zld (green, grayscale), Fas3 (red) and Tj (blue, grayscale).

(C) Graph showing relative Zld expression in the somatic cells of *tj*>+ (n=15) and *tj>Dcr-1-i* (n=15) testes.

Statistical analysis was performed using Student's t-test (**** P < 0.0001).

Scale bars: 20 μm.

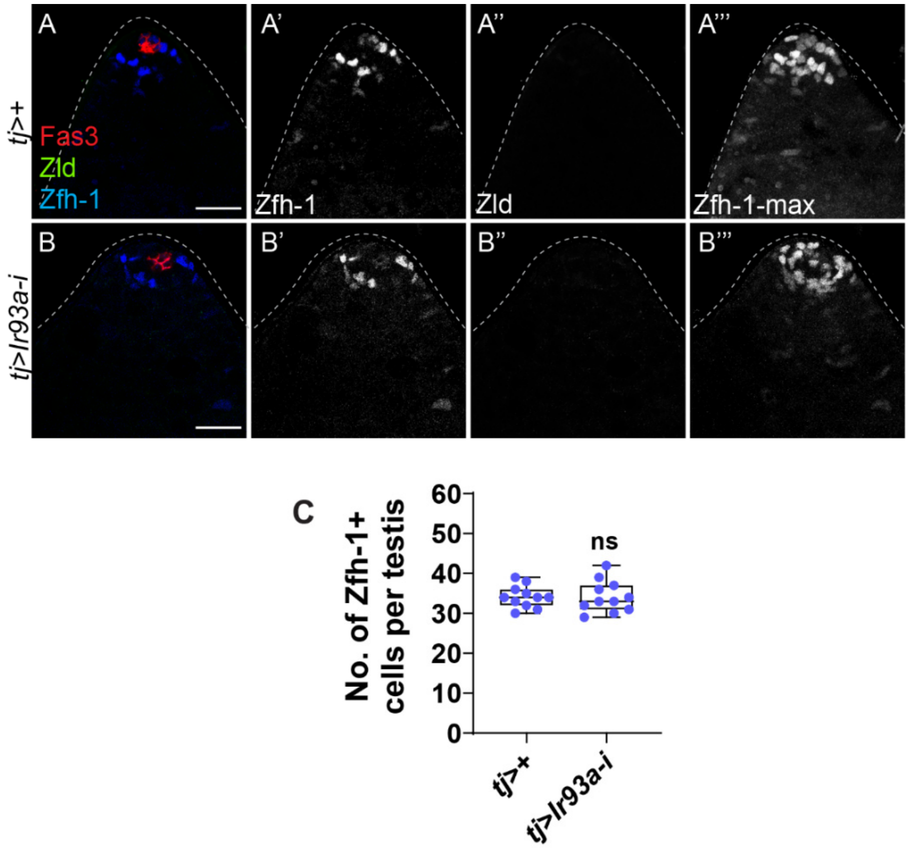

**Figure S5: Ir93a loss have no effect on CySC lineage, related to Fig. 3.**

(A, B) Representative confocal images of *tj*>+ (A) and *tj>Ir93a-i* (B) testes stained for Zld (green, grayscale), Zfh-1 (blue, grayscale) and Fas3 (red). A''' and B''' show Z-max projections of Zfh-1 expressing cells.

(C) Graph depicting the total number of Zfh-1 positive cells in *tj*>+ (n=11), *tj>Ir93a-i* (n=11) testes.

Statistical analysis was performed using Student's t-test (n.s. = not significant, P > 0.05).

Scale bars: 20 µm.

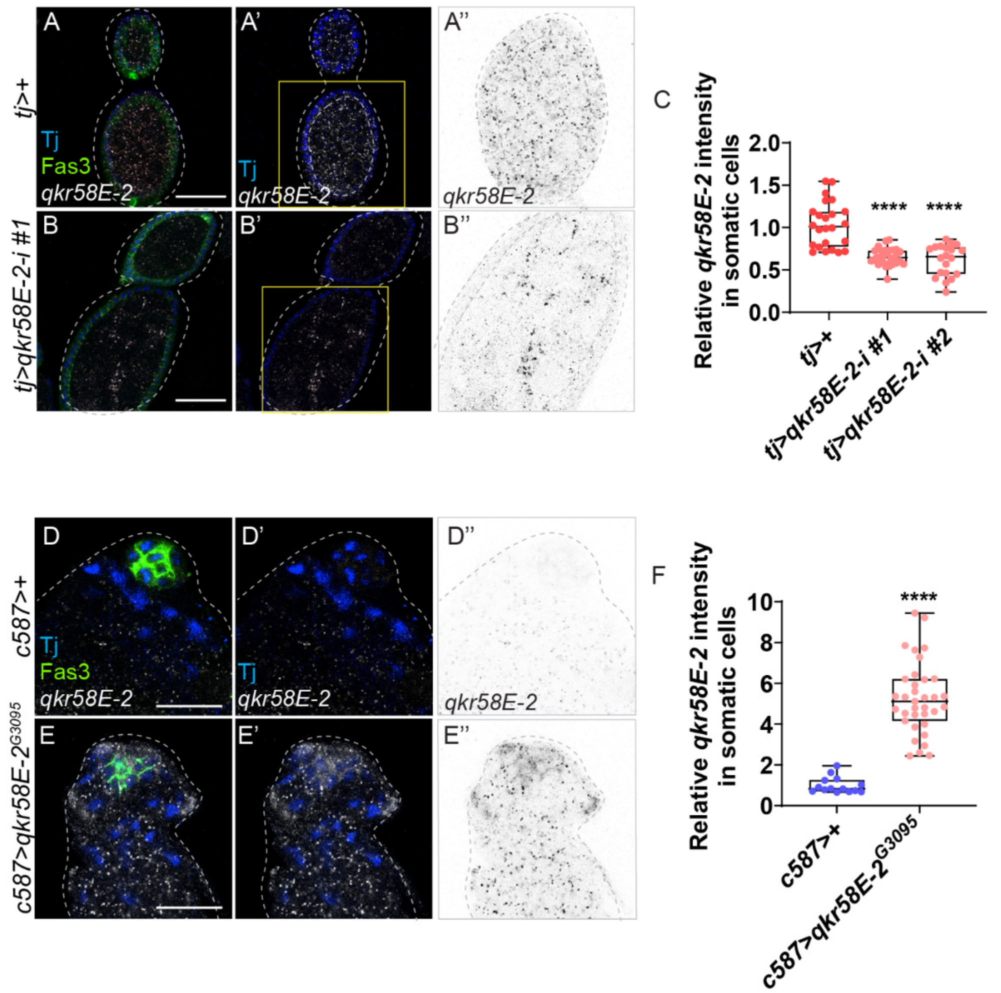

**Figure S6: Validation of *qkr58E-2* RNAi and *qkr58E-2$^{G3095}$ EP* lines, related to Fig. 5.**

(A, B) Representative confocal images of *in situ* hybridization for *qkr58E-2* mRNA in *tj*>+ (A), and *tj>qkr58E-2-i* (B) egg chambers. The egg chambers are stained for Fas3 (green), Tj (blue), and *qkr58E-2* mRNA (grayscale, inverted grayscale). The panels A" and B" shows the enlarged view of the inset marked with yellow boxes in A' and B'.

(C) Graph showing relative *qkr58E-2* mRNA intensity in the somatic cells in: *tj*>+ (n=8), and *tj>qkr58E-2-i* #1 (n=10) and *tj>qkr58E-2-i* #2 (n=8) ovaries.

(D, E) Representative confocal images of HCR FISH for *qkr58E-2* mRNA in *c587*>+ (D), and *c587>qkr58E-2$^{G3095}$* (E) testes. The testes are stained for Fas3 (green), Tj (blue), and *qkr58E-2* mRNA (grayscale, inverted grayscale).

(F) Graph showing relative *qkr58E-2* mRNA intensity in the somatic cells in: *c587*>+ (n=8), and *c587>qkr58E-2$^{G3095}$* (n=9) testes.

Statistical analysis was performed using Student's t-test (**** $P < 0.0001$).

Scale bars: 20 μm.

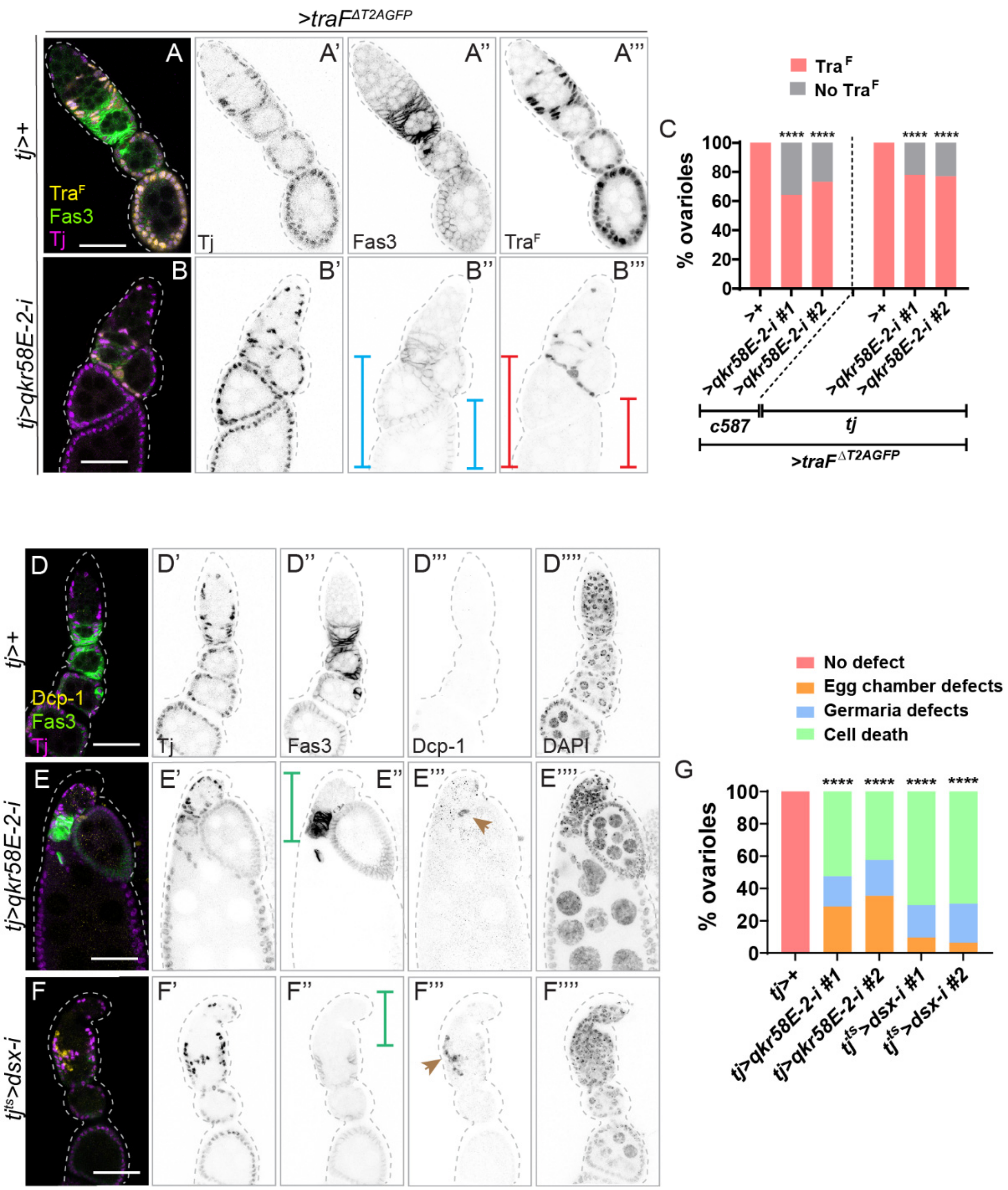

**Figure S7: *tra* alternative splicing and differentiation of ovarian follicle cells depends on *qkr58E-2*, related to Fig 5.**

(A, B) Representative confocal images of *tj> traF*$^{\Delta T2AGFP}$>+ (A), and *tj>traF*$^{\Delta T2AGFP}$>*qkr58E-2-i* ovarioles (B) stained for GFP (indicative of Tra$^{F}$) (yellow, inverted grayscale), Fas3 (green, grayscale) and Tj (magenta, inverted grayscale). Blue and red brackets in B'' and B''', respectively, show reduced Fas3 and Tra$^{F}$ in *qkr58E-2*-depleted follicle cells.

(C) Graph showing the percentage ovarioles with or without Tra$^{F}$ in *c587>traF*$^{\Delta T2AGFP}$>+ (n=65), *c587>traF*$^{\Delta T2AGFP}$>*qkr58E-2-i #1* (n=98), *c587>traF*$^{\Delta T2AGFP}$>*qkr58E-2-i #2* (n=71), *tj>traF*$^{\Delta T2AGFP}$>+(n=55), *tj>traF*$^{\Delta T2AGFP}$> *qkr58E-2-i #1* (n=68), *tj>traF*$^{\Delta T2AGFP}$>*qkr58E-2-i #2* (n=56).

(D-F) Representative confocal images of *tj >+* (D), *tj >qkr58E-2-i* (E), and *tj*$^{ts}$*>dsx-i* (F) ovarioles stained for Dcp-1 (yellow, inverted grayscale), Fas3 (green, grayscale), Tj (magenta, inverted grayscale) and DAPI (grayscale). Green brackets in E" and F" show fused egg chambers and abnormal germaria in follicle cells depleted for *qkr58E-2* (E") and *dsx* (F"). Brown arrows in E"' and F"' show Dcp-1 expression in follicle cells depleted of *qkr58E-2* (E"') and *dsx* (F"').

(G) Graph showing oogenesis defects and cell death in *tj* >+ (n=72), *tj >qkr58E-2-i #1* (n=52), *tj >qkr58E-2-i #2* (n=37), *tj*$^{ts}$*>dsx-i #1* (n=58) and *tj*$^{ts}$*>dsx-i #2* (n=32) ovarioles.

Statistical analysis was performed using Fisher's exact test (****$p < 0.0001$).

Scale bars: 20 μm.

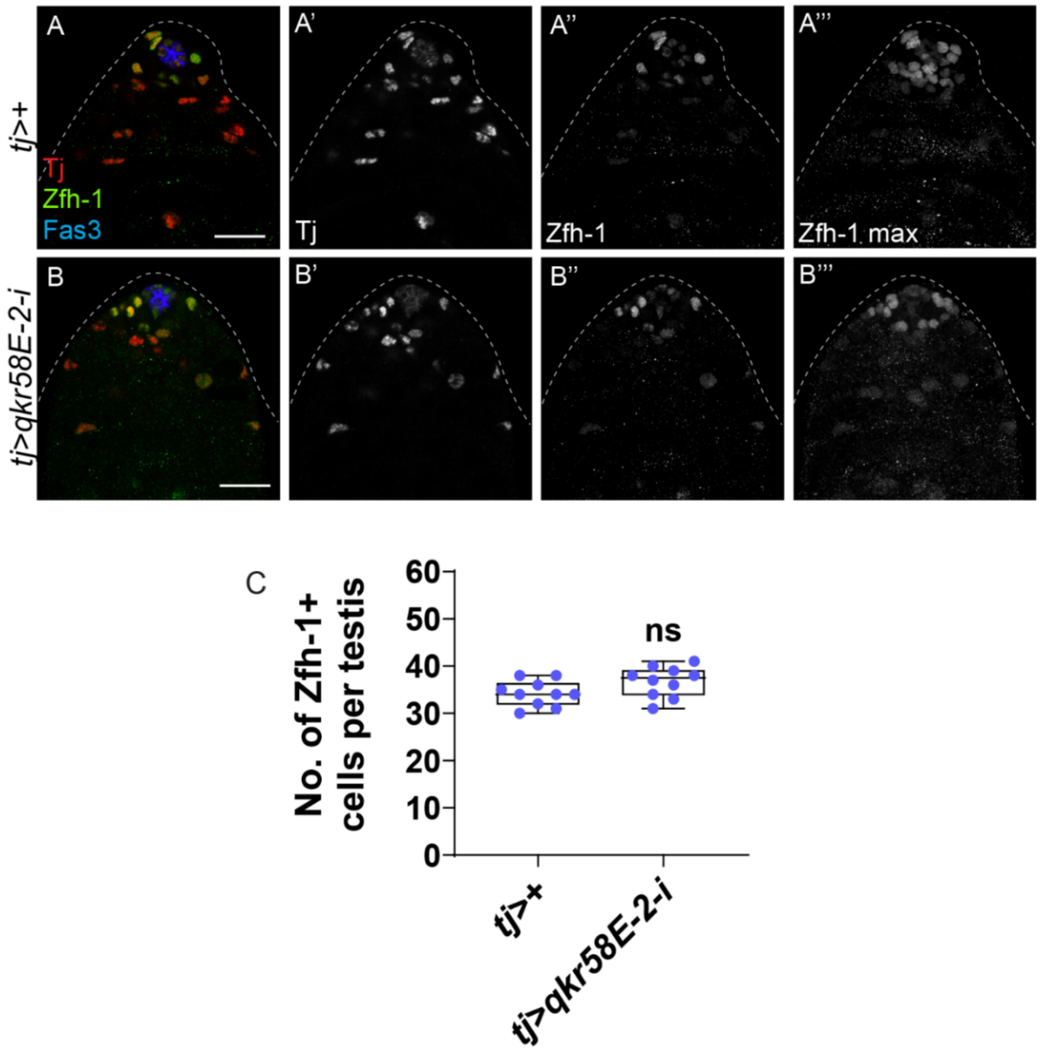

**Figure S8:** ***qkr58E-2*** **depletion does not impact the CySC lineage in the testis, related to Fig. 5.**

(A, B) Representative confocal images of *tj*>+ (A), and *tj>qkr58E-2-i* (B) testes. The testes are stained for Fas3 (blue), Tj (red, grayscale), and Zfh-1 (green, grayscale). A''' and B''' show Z-max projections of Zfh-1 expressing cells.

(C) Graph depicting the total number of Zfh-1 positive cells in *tj*>+ (n=10) and *tj>qkr58E-2-i* (n=10) testes.

Statistical analysis was performed using Student's t-test (n.s. = not significant, P > 0.05).

Scale bars: 20 µm.

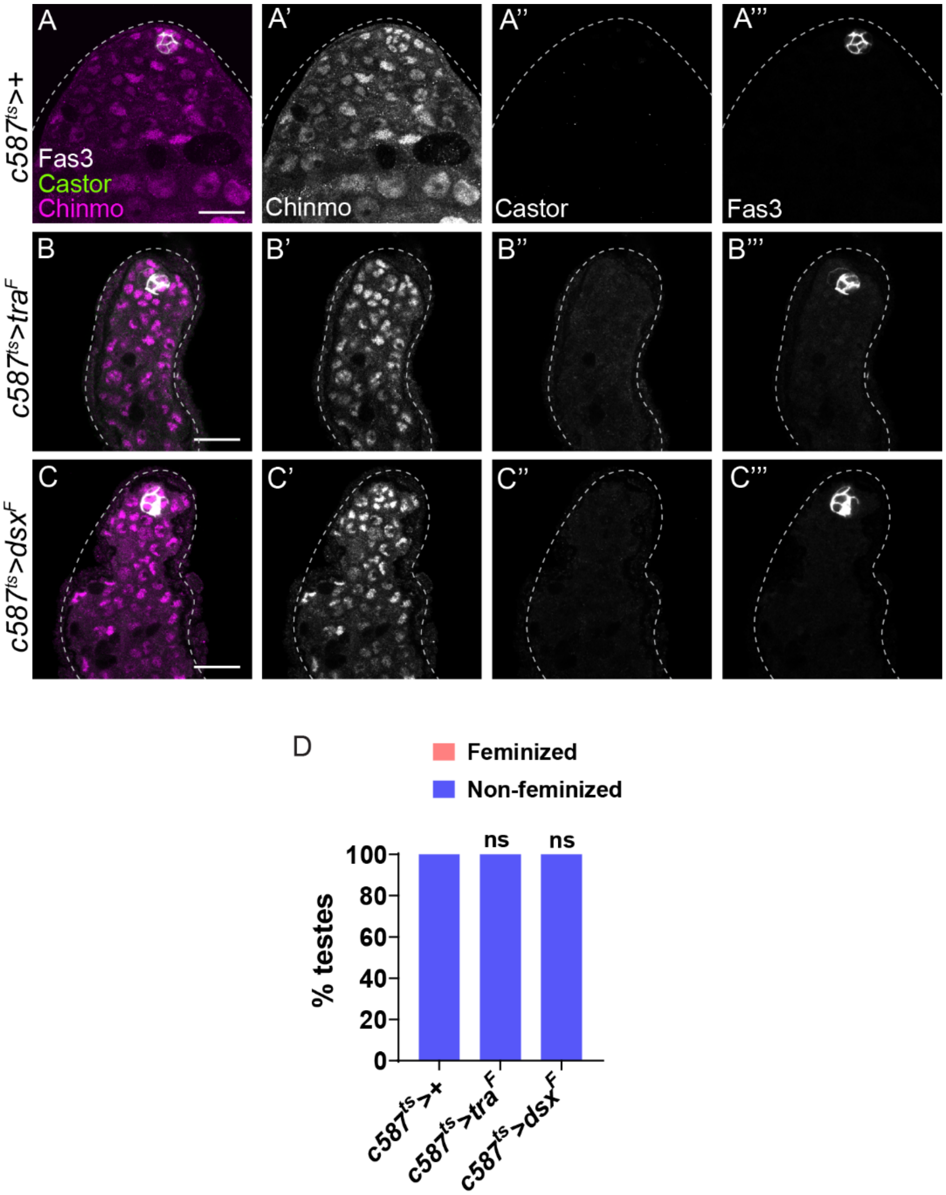

**Figure S9: Ectopic Tra$^{F}$ and Dsx$^{F}$ do not trigger Chinmo loss and feminization, related to Fig. 6.**

(A-C) Representative confocal images of *c587*$^{ts}$*>+* (A), *c587ts>tra*$^{F}$ (B), and *c587*$^{ts}$*>dsx*$^{F}$ testes at 20 days of adulthood. The testes are stained for Fas3 (gray, grayscale), Castor (green, grayscale), and Chinmo (magenta, grayscale).

(D) Graph showing the percentage of feminized and non-feminized testes in *c587*$^{ts}$*>+* (n=10), *c587*$^{ts}$*>tra*$^{F}$ (n=10) and *c587*$^{ts}$*>dsx*$^{F}$ (n=10).

Statistical analysis was performed using Fisher's exact test (n.s. = not significant).

Scale bars: 20 µm.

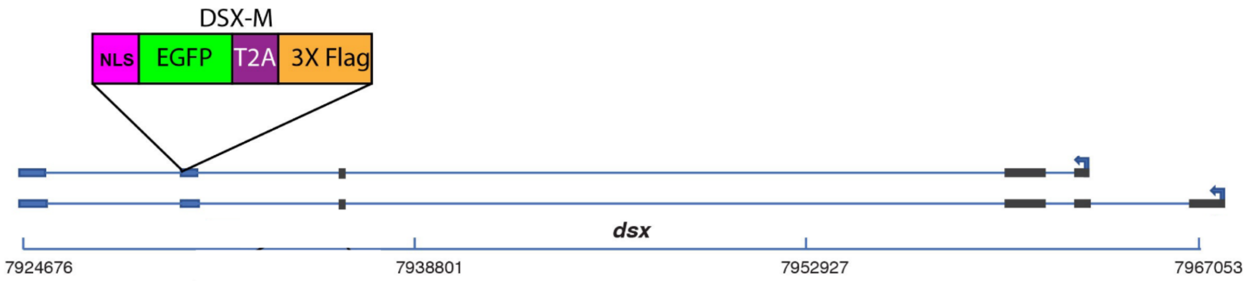

**Figure S10: Schematic for *DsxM::GFP allele*, related to Fig. 7.**

See Star Methods for detailed information.
